# Methods for constructing and evaluating consensus genomic interval sets

**DOI:** 10.1101/2023.08.03.551899

**Authors:** Julia Rymuza, Yuchen Sun, Guangtao Zheng, Nathan J. LeRoy, Maria Murach, Neil Phan, Aidong Zhang, Nathan C. Sheffield

## Abstract

The amount of genomic region data continues to increase. Integrating across diverse genomic region sets requires consensus regions, which enable comparing regions across experiments, but also by necessity lose precision in region definitions. We require methods to assess this loss of precision and build optimal consensus region sets. Here, we introduce the concept of *flexible intervals* and propose 3 novel methods for building consensus region sets, or universes: a coverage cutoff method, a likelihood method, and a Hidden Markov Model. We then propose 3 novel measures for evaluating how well a proposed universe fits a collection of region sets: a base-level overlap score, a region boundary distance score, and a likelihood score. We apply our methods and evaluation approaches to several collections of region sets and show how these methods can be used to evaluate fit of universes and build optimal universes. We describe scenarios where the common approach of merging regions to create consensus leads to undesirable outcomes and provide principled alternatives that provide interoperability of interval data while minimizing loss of resolution. Software is available at https://github.com/databio/geniml.

## Introduction

Advancements in high-throughput sequencing technologies have resulted in a vast amount of diverse epigenomic data that has given us tremendous insight into genome function. Epigenomic data are often summarized into genomic region sets stored in BED files. Through the work of hundreds of individual labs and projects such as ENCODE (1), the NCBI Gene Expression Omnibus (2) now contains almost 100,000 BED files (3). The volume of data has made integration challenging.

For an analysis that spans several genomic interval sets, one of the first steps is to define a consensus region set, or *region universe*, upon which the diverse sets can be interpreted (4–10). Such *universe* region sets have many common practical use cases. For example, they define genomic intervals for differential peak analysis(11); they form the regions of interest in a count matrix in single-cell epigenome analysis (11, 12); they are used as a background for statistical region enrichment analysis (13–16); and they are a region vocabulary in vector representation approaches (17–20).

For many tools, the choice of universe is critical. It defines the features to which data will be projected. Currently, there are different ways of choosing a universe for analysis. Simple approaches include tiling the genome into fixed-size bins (12, 15), or using intersection or union operations on a collection of region sets (11). Some methods have been developed to create better-fitting universes for specific downstream use cases (21, 22). An alternative is to use a predefined universe from an external source; for example, the ENCODE consortium curated a registry of candidate cis-regulatory elements accessible through the SCREEN webserver (1), and the Ensemble Regulatory Build is a central, reusable source of regulatory region definitions (23). The choice of universe matters because universes can be a poor fit to data, and if a universe does not fit the data well, it can lead to incomplete or incorrect results (5, 14, 15). However, despite the importance of selecting a universe, it is often done *ad hoc*, and there are few approaches to assess the fit of a universe to a collection of region sets.

Here, we address these limitations by introducing novel concepts for constructing and evaluating region universes. First, we introduce the idea of flexible genomic intervals, which represent region boundaries by intervals instead of points, allowing us to summarize many fixed regions into fewer flexible regions without loss of information. Next, we propose three methods for constructing flexible region universes: a coverage cutoff universe, a maximum likelihood universe, and a Hidden Markov Model (HMM). Finally, we propose three methods to evaluate the fit of a universe to a collection of region sets: 1) the base-level *F*_10_-score; 2) a Region Boundary Distance score (*RBD*); and 3) a likelihood model score that assesses the likelihood that the proposed universe was drawn from the given distribution of region sets.

To assess our universes and evaluation methods, we compared our methods against alternatives and predefined universes. We show that flexible universes can capture information from complex data collections into one well defined universe. Moreover, we show how our assessment metrics provide complementary measures of assessing universe fit and we prove the relevance of these measures. We show that the union universe has many downsides and propose the HMM universe as a generally useful approach for defining well-fit universes. To demonstrate how these universes could affect downstream analysis, we conclude with an application of region set enrichment analysis, where we show how the results are affected by choice of universe. Overall, our results demonstrate the importance of considering region universe and provide promising new tools to construct better-fitting universes for a variety of use cases.

## Methods

### Overview

To integrate genomic interval data, we first require a consensus set of intervals, or a universe. We may select a predefined universe from an external source or define one from a collection of input region sets using a consensus algorithm (Fig. 1A). With a universe in hand, we can then “tokenize” the original regions (Fig. 1B). Tokenization redefines them into universe regions, normalizing differences in region boundaries to transform similar regions into a single representation. The most basic tokenization method is simple interval overlap. This approach works well if a universe approximates the original data well; otherwise, this may result in loss of precision. A universe may not be a good fit to a collection for a variety of reasons (Fig. 1C); for example, 1) a region can be shifted; 2) two neighboring regions may be merged, making them indistinguishable in downstream analysis (5); 3) a universe may omit important intervals, leading to loss of information (5); or 4) a universe may contain extraneous regions that do not reflect genome coverage, adding noise and compute time (14). If a universe is a poor fit, it can affect downstream analysis negatively; for example, a differential accessibility analysis wouldn’t even test a locus that had been dropped from the universe. It could also miss a significant locus if it were merged with an abutting locus that lacked differential signal. Or, a motif analysis based on universe regions that had been shifted could miss enriched sequence features that were present in the part of the region left out of the universe.

**Figure 1:**
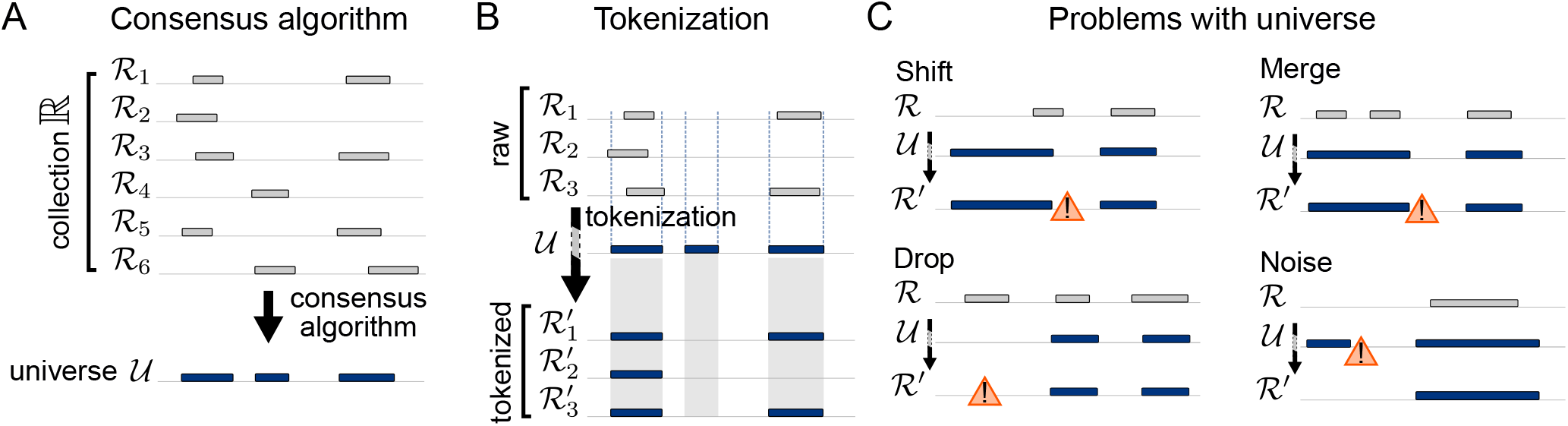
Overview of the concept of a universe. A) A consensus algorithm takes a region set collection ℝ = [ ℛ_1_, ℛ_2_, …] and builds a consensus representation called universe 𝒰. B) We “tokenize” raw regions into universe 𝒰 by redefining them as universe regions, creating a more uniform collection. C) Universes may poorly represent region sets by shifting, merging, dropping, or adding extraneous regions.

To address these issues, we developed three new approaches for constructing a universe that is a good fit to the original data: first the “coverage cutoff universe”; second, the “maximum likelihood universe”; and finally, an “HMM universe”. To assess them, we also developed three universe fit metrics. Finally, we applied these to a variety of real datasets.

### Methods for building optimal universes

#### The coverage cutoff universe

The simplest example of a universe built from the data is a “union universe”, in which a collection of region sets is merged. This method is often done for differential analysis of ATAC-Seq data (11). The union universe by definition covers all bases from the original collection; however, it can also lead to very large regions, particularly if the number of input region sets is large. Another simple alternative is using an intersection operation, which would include only bases covered in every region set in the collection, but this has the opposite problem: it leads to very sparse universes.

We reasoned that a hybrid approach may achieve a better result. First, we conceptualize a collection of region sets as a coverage signal track across all input region sets. Then, similar to a peak calling approach, we choose a cutoff *x* such that universe includes only positions with coverage greater or equal to *x* (Fig. 2A). Setting the cut-off to one corresponds to a union universe, and setting the cutoff equal to the number of input region sets corresponds to an intersection universe. Setting a cutoff in between the two balances these extremes and provides a tunable parameter that may be adjusted depending on the needs of downstream tasks. We call the resulting universe a coverage cutoff (CC) universe. A principled approach to selecting the cutoff is to use a simple likelihood model that calculates the probability of appearing in a collection. With this model, we can calculate an optimal cutoff according to Eq. 1 (see Supplementary methods for details).

**Figure 2:**
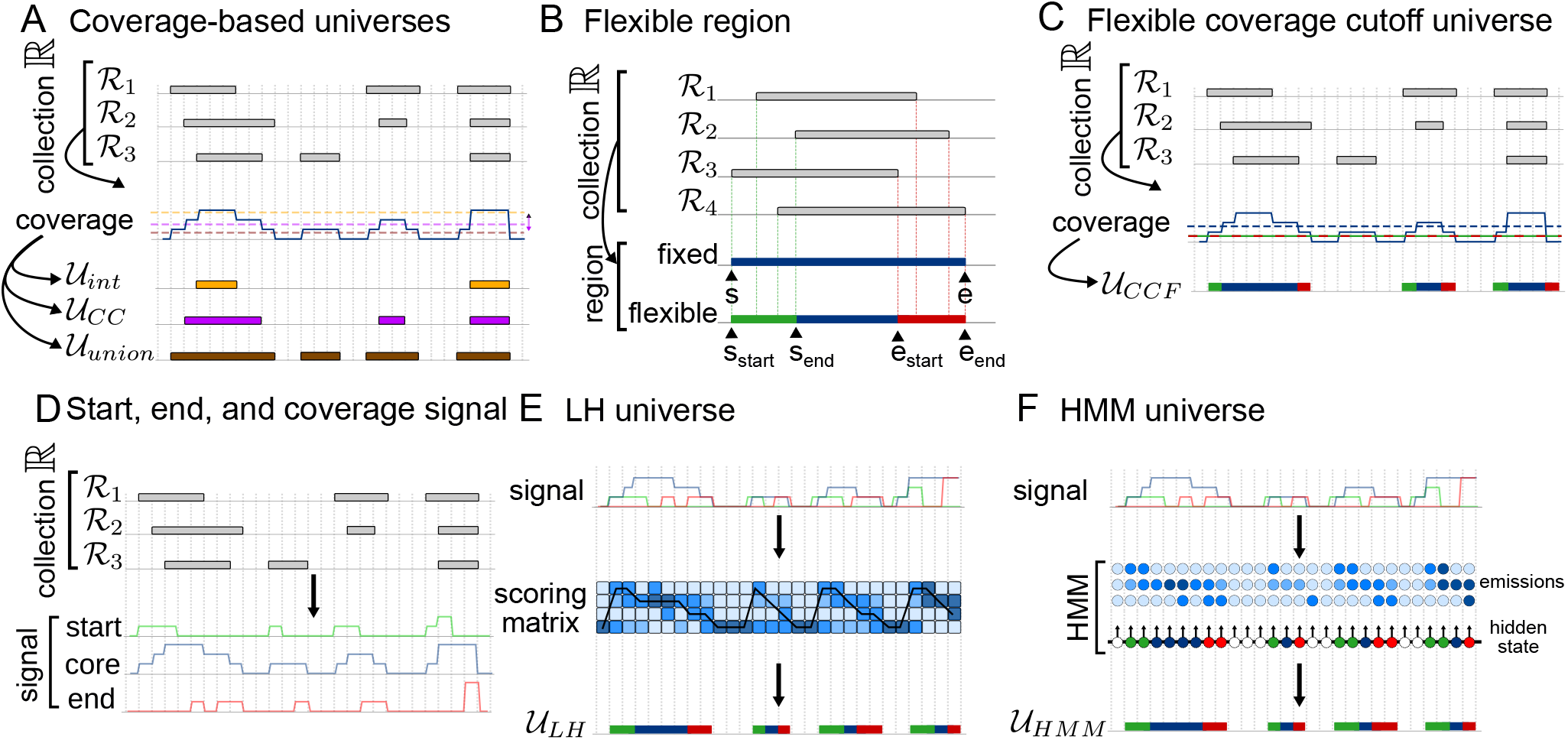
Different approaches to building universes. A) Coverage-based universes are derived from the genome coverage of a collection of region sets. Examples include intersection 𝒰_int_, coverage cutoff 𝒰_CC_, and union universe 𝒰_union_. B) A flexible region in contrast to fixed region can represent boundaries of many variable regions. C) The flexible coverage cutoff (CCF) universe is based on coverage of the genome by a collection. It uses two cutoff values: the lower defines flexible boundaries and the upper defines the region core. D) A collection of genomic region sets is aggregated, and region starts, core (overlap), and ends are counted, creating signal tracks. E) Maximum likelihood universe is derived from three signal tracks. Using a likelihood model, we build a scoring matrix that assesses the probability of each position being a given part of a flexible region. Next, we find the most likely path, which represents the maximum likelihood universe. F) The HMM universe treats signal tracks representing genome coverage by different parts of a region as emissions of hidden states that correspond to different parts of flexible regions.

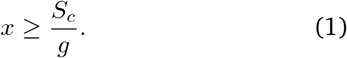

Here, *S*_*c*_ is a sum of genome coverage by collection and *g* is the size of the genome.

We realized that, in a sense, the CC universe is a point estimate of a more complex distribution of possible universes. We reasoned that we may gain some insight by modeling the boundaries of the consensus regions as intervals, rather than points. To do this, we developed a new concept of a genomic region we call a *flexible region*. In contrast to fixed interval that is defined by two fixed points (*start* and *end*), a flexible interval is defined by four (*start*_*start*_, *start*_*end*_, *end*_*start*_ and *end*_*end*_) (Fig. 2B). A flexible interval can model many region variations into one well-defined flexible interval.

A simple approach to constructing a flexible region universe is to define two cutoff values instead of one: the looser represents the cutoff for boundaries, and the stricter for the region core (Fig. 2C). This way, positions with coverage between those two points will be assigned to flexible region boundaries and positions with coverage higher than the second cutoff will be assigned to core of the region. Using this idea, we built a confidence interval around the optimal cutoff value, which extends the CC universe into the coverage cutoff flexible (CCF) universe.

#### Maximum likelihood universe

While flexible intervals more naturally represent collections of overlapping region sets than fixed intervals, we reasoned that they still suffer from the possibility of merging neighboring regions when collections are large. To address this issue, we need information about not just the coverage of regions in a collection, but also the region start and end positions. To compute this information, we developed a fast plane-sweep algorithm (see Supplementary methods for details). With this tool we can quickly and efficiently calculate three tracks representing aggregate start, end, and coverage values of a region set collection at base-pair resolution (Fig. 2D). We reasoned that a model that could incorporate all of these signals may improve universe resolution.

We can conceptualize a flexible universe as a path through the genome that assigns either start, core, or end state to each position. Using a universe scoring model and optimization algorithm, we can build the best path through the genome (Fig. 2E). As a scoring model, we next developed a complex likelihood model, an extension of the simple likelihood model introduced earlier for CC universe, which considers not just the coverage (core), but also in region start and end signal tracks. This model describes for each position the probability of it being a region start, core, or end. We thus build the *likelihood universe* (LH) in 3 steps: 1) compute the three signal tracks; 2) use a likelihood model to build a scoring matrix; 3) find the maximum likelihood path through the genome (see Supplementary methods for details).

#### Hidden Markov Model universe

The maximum likelihood universe provides a simple and principled model for optimal flexible universes. However, a disadvantage is that it provides no tunability, since the likelihood scores are determined purely from the data. We reasoned that this may lead to results depending on input collection. To address this, we sought a more tunable model using a Hidden Markov Model (HMM).

An HMM models a hidden processes using 1) a matrix of transition probabilities between hidden states; and 2) emission probabilities of observations from hidden states. In our model, there are three observed sequences: the number of starts, overlaps, and ends at a given position. The hidden variable corresponds to the different parts of the flexible segment (Fig. 2F; S1). We can tune transition probabilities, which can be chosen in a way that will prevent unnecessary segmentation, and emission matrix, which describes the relationship between observations and hidden states (see Supplementary methods for details).

### Methods for evaluating universe fit

Having developed several new approaches to construct universes, we next sought to evaluate these universes and compare them to other common approaches. Because the choice of universe can dramatically affect downstream analyses, it is important to choose a universe deliberately. However, there are no well-established methods for assessing universe fit to data. Furthermore, different analyses may be better served by a different types of universe, indicating that there really is no generally optimal universe, but the idea of what makes a “good” universe depends on the downstream analysis. For example, a differential accessibility analysis should prioritize sensitivity over specificity; in this case, it is not a major problem if the universe includes many regions present in only a few samples, since the cost of extra comparisons is lower. On the other hand, the cost of excluding a region that could be a significant differential locus would be high. In contrast, a word-based deep learning task that trains a model with input dimensions equal to the size of the universe may elect a more specific universe, at the cost of discarding some regions that are present in a few of the samples, because otherwise the training could be intractable. Thus, the question of universe optimality depends on the use case, and therefore, we require methods of evaluating universe fit that can be tuned to a research question. This problem is similar to the comparison of two generic interval sets, for which several methods have been developed (24), with two key differences: first, we want to compare a universe region set not only to one other region set, but to a collection of them; and second, the question is not symmetric: it is generally more important that a universe not miss information (regions), even at the cost of some extra regions – and the desired level of asymmetry can vary. Therefore, we developed three methods for assessing universe fit to data: 1) a base-level overlap score; 2) a region boundary distance score; and 3) the universe likelihood.

#### Base-level overlap score

Our first metric is based on base coverage. We consider the universe as a prediction of whether a given genomic position is present in a given region set from the collection. Treating each region set from the collection as a query, we can then conceptualize matches and mismatches as true positives (covered in both universe and query), false positives (covered in universe, but not in query), or false negatives (covered in query, but not in universe) (Fig. 3A). This allows us to calculate common classification evaluation measures such as precision and recall. Precision counts the number of true positives, so a low precision indicates presence of unimportant positions in the universe; recall measures how much of the universe is in a query. To combine precision and recall, we use the *F*_10_-score, a weighted version of the traditional *F* -score that pays 10 more times attention to recall than precision. This asymmetry captures our goal to prioritize sensitivity over specificity: by prioritizing recall, we indicate that it’s better to have a few extra, noisy regions that to exclude something important. The *F*_10_-score results in values between 0 and 1, with a perfect fit approaching 1. An alternative approach to baselevel overlap score would be Jaccard similarity, however it does not account for asymmetry of the comparison.

**Figure 3:**
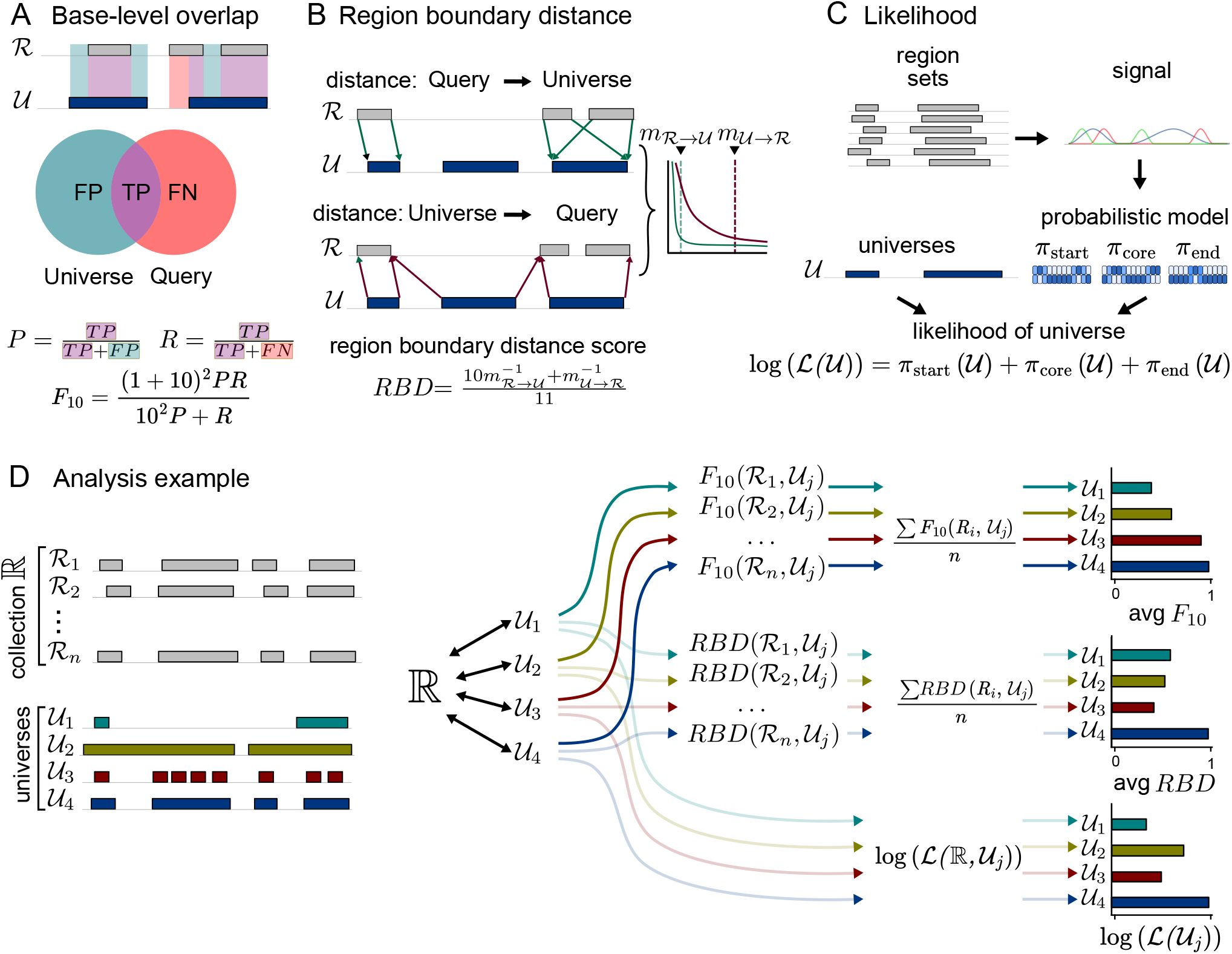
Different approaches to assess how well the universe represents the data. A) Base-level overlap measure considers universe a prediction of a region set and based on that it calculates number of false positives (FP), true positives (TP), and false negatives (FN), and from that derives recall (R) and precision (P), which are combined into F_10_-score. B) Region boundary distance assesses how well a universe represents start and end positions, by calculating distance from region set to universe, and from universe to region set; those two metrics are combined into a region boundary score by calculating their reciprocal, weighted harmonic mean. C) Likelihood assessment uses a likelihood model based on signal tracks representing genome coverage by different parts of a region to calculate universe likelihood as a combination of likelihoods of all three signals tracks. D) A complete analysis example comparing a collection of region sets against 4 proposed universes: 𝒰_1_ a precise universe, 𝒰_2_ a sensitive universe, 𝒰_3_ a fragmented universe, and 𝒰_4_ well-fit universe. For each universe, all 3 metrics are calculated. The F_10_-score and RBD score assess individual region sets. The final score for a collection is their average. In contrast, the likelihood is calculated directly for the whole collection.

#### Region boundary distance score

One disadvantage of the base overlap score is that it is unaware of region boundaries. A universe region that covers two abutting query regions would get a perfect score. This can be highly problematic in downstream applications; for example, in a differential analysis, lumping two distinct loci together could dilute differential signal. To address this, we sought a measure that would consider region starts and ends (Fig. 3B). We calculate the distance between each boundary of each region in the query and the closest corresponding boundary in the universe. Universes with boundaries that are near the query boundaries would have shorter distances, indicating better universe fit. However, highly fragmented universes with many unnecessary boundaries would have very small distances from query to universe. To account for this, we also calculate the inverse distance: from boundaries in universe to the nearest boundaries in the query. Finally, we combine those two metrics into region boundary distance score (*RBD*) by taking their reciprocal, weighted harmonic means. With this score we describe the universe’s ability to conserve information about starts and ends, with one representing a perfect representation of boundary locations. However, we do not incorporate any information about collection coverage in this score.

For fixed universes, the start and end point are well defined, however for flexible regions they are intervals. Therefore, for flexible universes, we modify the *RBD* score to set distance equal to zero if a boundary query region is inside the universe’s boundary interval.

#### Universe likelihood

Finally, we sought a metric that incorporates both information about region boundaries as well as genome coverage. We propose here a universe likelihood score (Fig. 3C). We first calculate three signal tracks representing genome coverage by start, core and end of the regions in the collection. Then for each signal track we make a separate model, which results in three separate models for different parts of a region. Each of these models describes the probability of a given position being a given part of a region, depending on the signal strength (see Supplementary methods for details). That results in a complex, probabilistic description of a region set collection. Next, we use this model to calculate the likelihood of the universe, which we can compare between universes. We make two versions of likelihood calculations, one suited for fixed universes and one for flexible universes. We use log likelihood, so our values range from minus infinity (low) to zero (high). Finally, to increase interpretability, we normalize the scores by subtracting the likelihood of an empty universe (one that contains no regions). Thus, a positive final score reflects a given universe that is more likely than having no regions at all, while a negative score means the universe is less likely than the empty universe.

To accommodate flexible universes, we adapted the likelihood score by calculating the boundary likelihood as if the whole flexible interval could contain a boundary position, rather than a single fixed point (see Supplementary methods for details).

#### Assessing region sets collections

Having developed 3 assessment methods, we can use them to compare competing universes to assess which universe is the best fit for a collection of region sets. We do this by computing the scores for each universe and comparing among universes (Fig. 3D). The scores assess different aspects of the universe fit; the *F*_10_-score promotes sensitive universes over specific ones, *RBD* score penalizes sparse universes, and likelihood provides complex universe assessment. Although the likelihood score incorporates information about boundaries as well as how well the universe covers the collection, it can penalize sensitive universes because it is not intentionally biased toward asymmetry the way the previous scores are. Thus, by computing all these scores, we reason that we get a complete picture that can guide decisions for selecting a universe for a collection of region sets.

### Evaluation on real data

Next, we developed an evaluation strategy to test our universe building and assessment methods on real data. We assembled five diverse collections of region sets representing different biological problems (Fig. 4A): 1) *CTCF ChIP small*, a small random collection of CTCF region sets (n=40) from the ENCODE database (1); 2) *CTCF ChIP large*, CTCF ChIP-seq datasets (n=877); 3) *TF ChIP*, ChIP-seq experiments for diverse transcription factors (TFs) (n=8,503); 4) *B-LCL ATAC*, a small set of ATAC-seq files from B-Lymphoblastoid cell lines (B-LCL; n=400) from ChIP-Atlas (25); 5) a *Random ATAC*, random ATAC-seq results (n=5,000). These datasets vary in data type, collection size, and level of heterogeneity of input regions across region sets.

**Figure 4:**
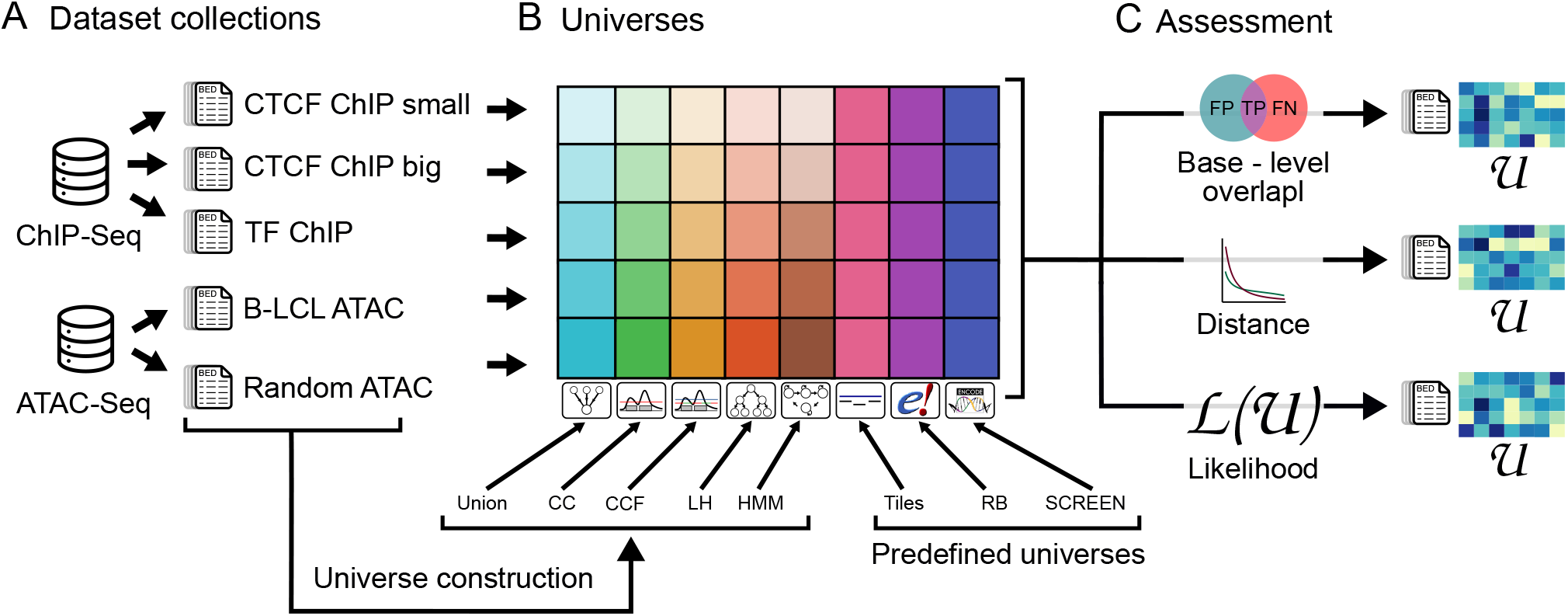
Overview of evaluation approach. A) Five collections representing different biological problems used for assessment. B) For each collection, we compared it to five data-driven universes and three predefined universes. The data-driven universes are tailored to the input collection, but the predefined universes do not vary by collection. C) We assessed the fit of each universe to each collection using our 3 assessment methods.

We also assembled universes to assess. First, we obtained 3 universes that do not depend on analyzed data, which we call *predefined* universes: 1) the tiles universe, which bins the genome into non-overlapping 1000 bp tiles; 2) the SCREEN universe, which consists of predefined cis-regulatory elements from ENCODE (1); and 3) the Regulatory Build (RB) universe, consisting of pre-defined regulatory elements from Ensembl (23). In addition to these 3 external universes, we also built 5 data-driven universes that are specific to each region set collection. These include: 1) the union universe; 2) the CC universe; 3) the CCF universe; 4) the LH universe; and 5) the Hidden Markov Model (HMM) universe. This led to 28 universes and 40 pairwise comparisons of universe-to-collection (Fig. 4B). For each comparison, we computed our three assessment methods (Fig. 4C). This gives us a comprehensive evaluation of both externally sourced and data-driven universes, tested on diverse query region set collections.

## Results

### Data overview

To explore the differences in our five region set collections, we first computed general coverage statistics (Table S1). The smallest region set, *B-LCL ATAC* contains ≈ 700,000 regions and covers 0.2% of the genome, whereas the largest, *TF ChIP*, contains ≈ 1.5 billion regions and covers 91% of genome. We also observed that the ATAC-seq collections have smaller regions on average than the ChIP-seq collections.

### Universe overview

The universes also have very different characteristics, with some requiring additional filtering based on region likelihood and size (see Supplementary methods for details, Fig. S2, S3). For example, for the *CTCF ChIP large* collection, the 8 universes have different levels of precision and fragmentation (Fig. 5A). The universes differed in average region size, number of regions, and percent of genome covered (Fig. S4, Table S2). Having assembled the universes, we next computed our 3 assessment methods.

**Figure 5:**
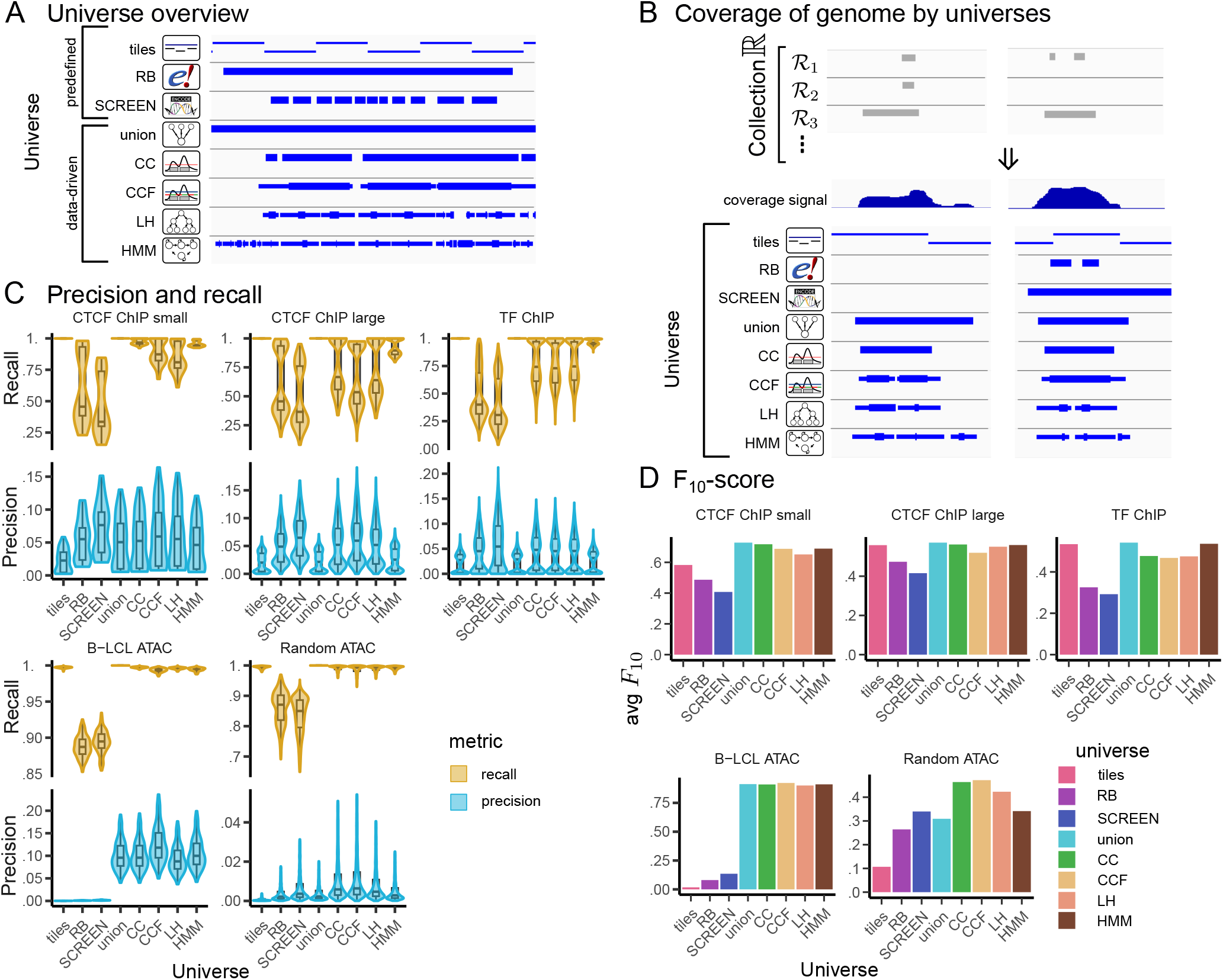
Universes overview and results of base-level overlap score. A) Example of universes assessed for the CTCF ChIP large collection, including the 3 constant external universes, and 5 data-driven universes built from the input collection. B) Different universes represent genome coverage by the collection to a different extent, example from the Random ATAC collection. Collection ℝ consists of many different files, which represented by core signal track. Regions in ℛ_1_, ℛ_2_, ℛ_3_ are best represented by CC, CCF and LH universes in terms of overlap. C) Precision and recall distribution for each collection and universes assessments. D) Average F_10_-score for each collection and universes assessments

### Assessment 1: base-level overlap *F*_10_-score

Region sets in a collection can differ widely; our first assessment method assesses fit by quantifying the degree of overlap between each region set and the universe. In example data from the *ATAC Random* collection, we observe that some universes cover many bases present in only few of the collection’s region sets, while other universes are more stringent (Fig. 5B). To assess this globally, we first computed precision and recall for each comparison (Fig. 5C). We observed that the tiling universe and union universe both have perfect recall for all tested collections, consistent with how these universes are constructed; the tiling universe covers the entire (mappable) genome, and the union universe by definition covers every base contained in the collection. In contrast, recall is lower for the more stringent datadriven universes; the CC, CCF and LH universes exclude positions with low coverage, especially for large collections; the HMM universe has higher recall in general, indicating that it contains most positions covered by the collection. Finally, the lowest recall scores are assigned to the external universes, SCREEN and RB, which is consistent with these universes being built from other data sources. This highlights an advantage of building bespoke universes tailored to a collection: recall is superior. On the precision side, the worst performer overall is the tiles universe, consistent with many tiles in the universe that do not reflect coverage in the collection. In contrast, SCREEN and RB had generally higher precision, especially for ChIP-Seq collections.

In general, data-driven universes tend to represent collections well with good precision and recall. This is most apparent for the *B-LCL* collection for which datadriven universes are much better than predefined. For likelihood universes (CC, CCF, and LH universes) built from large ChIP-Seq collections (*CTCF ChIP large* and *TF ChIP*) we observe the worst recall among data-driven universes. This is the consequence of Eq. 1, from which we observed that, for ChIP-Seq collections, the optimal cutoff value is higher. On the other hand, both the HMM universe and the union universe have high recall and low precision. In general, we see that universes with high recall have lower precision, reflecting the delicate balance between including complete information in universes without adding too much noise.

To propose a balance between precision and recall, we next calculated the *F*_10_-score, which assigns more weight to recall than precision (Fig. 5D). For predefined universes, we see that the tiles universe scores well for ChIP-seq collections, which cover more of the genome, but poorly for ATAC-seq collections, which have lower coverage. In general, both RB and SCREEN are outperformed by data-driven universes, especially for the *B-LCL ATAC* collection, for which they contain too much noise. In general, for ChIP-Seq collections, the union universe is the best for these metrics, consistent with the weighting we chose that gives 10 times the weight to recall. Interestingly, for the *TF ChIP* collection, the HMM universe outperforms likelihood universes; on the other hand, for ATAC-Seq collections, the CCF universe outperforms both union and HMM universes. Overall, we conclude that computing precision, recall, and *F*_10_ provide useful insight into assessing universe fit. They provide a way to quantify the advantages of a data-driven universe and assess how much information is lost by an external universe.

### Assessment 2: region boundary distance score

Next, we sought to address the major weakness we see in the base-level overlap score: that it does not consider region boundaries. Assessing boundaries is important because of how it affects downstream analysis. If two regulatory elements with distinct behavior are merged into a single region in a universe, then all downstream analyses will essentially evaluate the average of the two signals. But different universes have different sensitivities to boundary points (Fig. 6A); for example, anecdotally, the union universe is not sensitive at all: it contains few boundaries, especially for larger collections. It will clearly merge together many neighboring regions, even if they have distinct patterns across input sets. The CC and CCF universes are more sensitive but still miss out on many boundaries for bigger collections; the LH universe is very sensitive to boundaries, but has other weaknesses (it tends to exclude positions with low coverage); all of those problem are solved with HMM universe, which is very sensitive and also is able to represent regions with low coverage. To assess boundaries globally, we turned to the region boundary distance score. First, for each region set, for each region, for each boundary, we calculated the distance to the nearest corresponding universe boundary (See Methods; Fig. 6B). For all collections, the distance from collection to RB universe was very high. Similarly, the union universe performs very poorly in this metric, particularly for larger collections, consistent with intuition that the union re-gions lose boundary precision as the number of regions increases. We also computed the inverse: distances from query to universe (Fig. 6C). Interestingly, for ATAC-Seq collections, we observe a small distance from query to universe but a high distance from universe to query. This indicates that all universes have many boundaries that are not present in the raw data, but at the same time boundaries present in the queries are well-reflected by universes.

**Figure 6:**
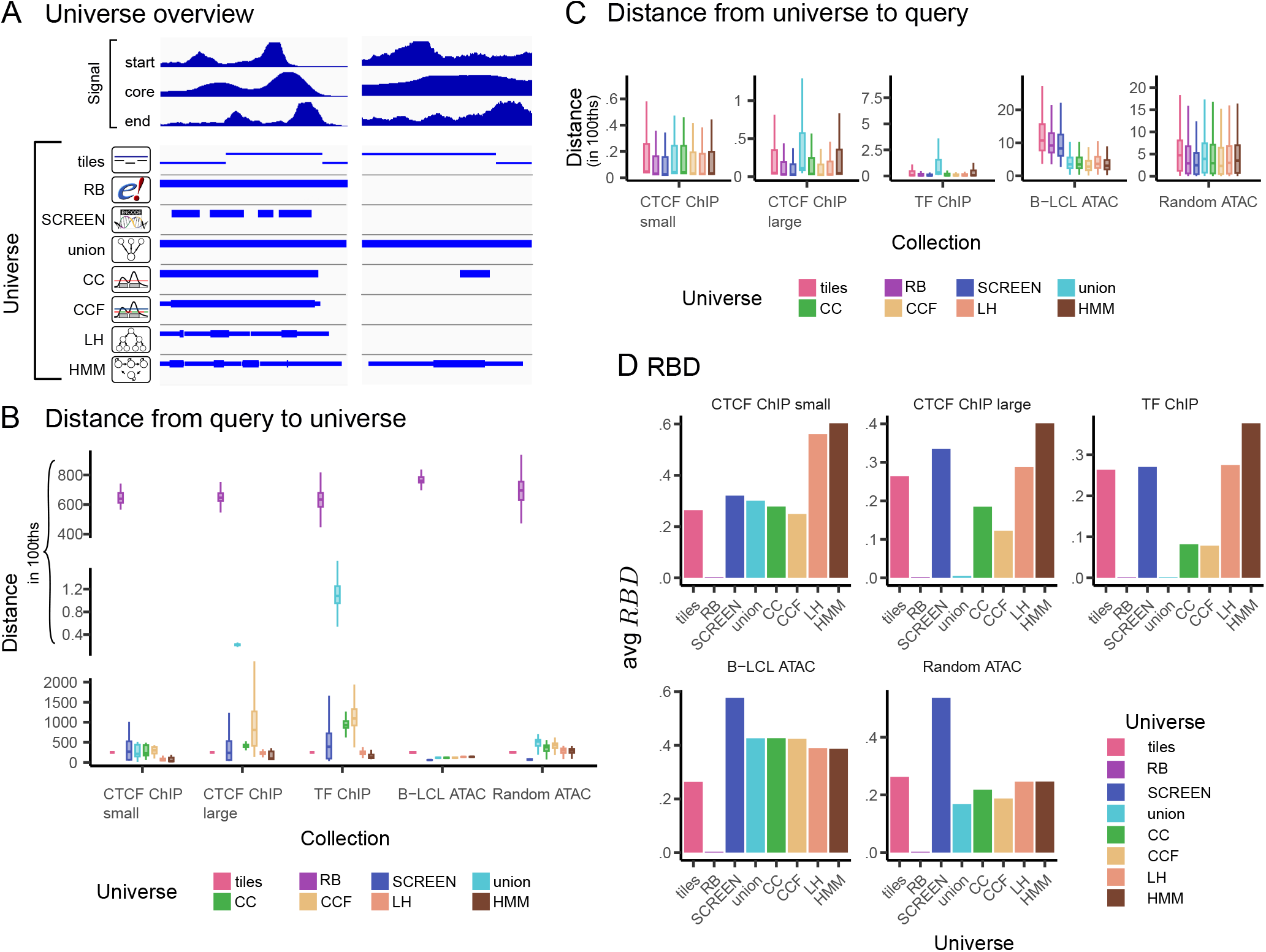
Results of region boundary distance score. A) Different universes represent region boundaries to a different extent. Three signal tracks provide summarized description of the whole collection. Both LH and HMM universe are most sensitive to region boundaries. B) Distribution of median distances from query to universe. C) Distribution of median of distance from universe to query. D) Average RBD score for each collection and universe comparison.

To summarize both distance directions, we calculated their reciprocal, weighted harmonic mean, the Region Boundary Distance score (*RBD*) (Fig. 6D). The average *RBD* score shows that the RB universe is a very poor fit to all collections, likely because it contains few, large regions. On the other hand, tiles universe has similar scores for all collections, which is good for big ChIP-Seq collections compared to other universes, but bad for ATAC-Seq collections. Interestingly, the SCREEN universe seems to be a good fit for all collections, and the best fit for ATAC-Seq collections, even outperforming the data-driven ones for this metric. Among datadriven universes, the HMM universe performs the best, with LH in second place for all collections except *B-LCL ATAC*. However, for this collection all data-driven universes have similar scores. Coverage-based universes (CC, CCF) perform well for ATAC-Seq collections, but not for large ChIP-Seq collections compared to other datadriven universes. As expected, the *RBD* score reflects the poor performance of the union universe for large ChIP-Seq collections; the merging leads to poor reflection of interval boundaries.

### Assessment 3: likelihood calculation

Finally, we calculated the likelihood score for each comparison (Fig. 7A). In general, predefined universes are a worse fit to the collections than an empty universe, with exception of SCREEN for *CTCF ChIP large* and *TF ChIP* collections. Among data-driven universes, the union universe performs very poorly, achieving negative scores for all collections. As expected, CC, CCF, and LH universes outperform the empty universe for all collections, as these were designed to optimize likelihood in some way. The HMM universe performs well overall; however, it is worse than empty universe for *CTCF ChIP large* and *TF ChIP* collections. A more detailed look reveals that the low HMM likelihood scores in these scenarios are driven by region coverage tracks, not boundaries, suggesting that for these collections, our current HMM parameterization may yield a universe with too much noise (see Methods; Fig. S5).

**Figure 7:**
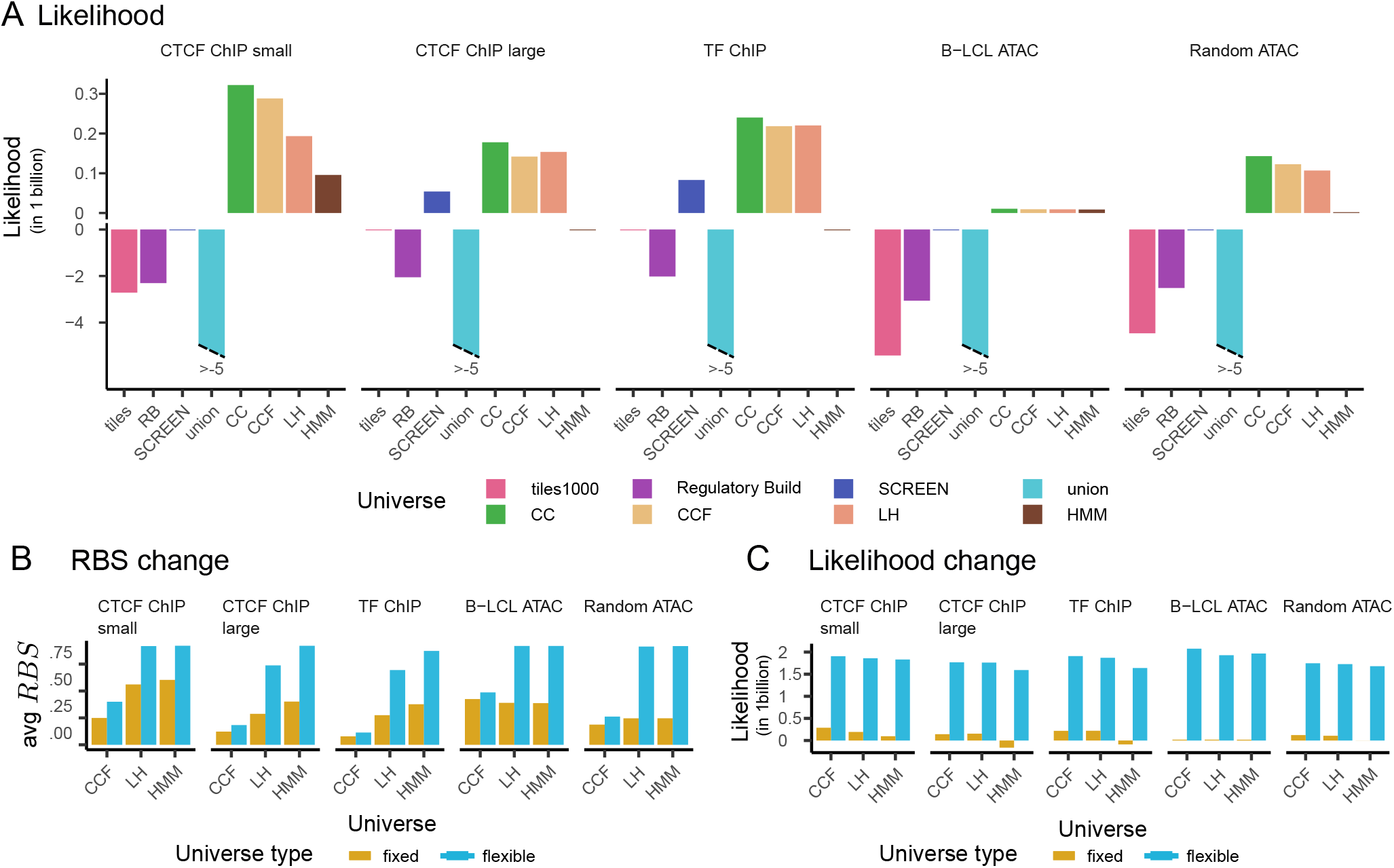
Results of universe likelihood and comparison between fixed and flexible scores. A) Likelihood of each universe given collection. B) Change of RBD score when we account for flexibility. C) Change of likelihood, when we account for flexibility.

### Comparing flexible to fixed universes

So far, our assessments have not taken into account that some universes can be flexible. We believe that flexible universes provide several advantages, and sought to assess them. Flexible intervals don’t quite fit into the standard 3-column BED format; however, they can be stored using optional fields of an extended BED format. In this approach, sequence name, start, and end of the flexible region are represented by three first columns, and thickStart and thickEnd columns hold the information about end of the flexible start and start of the flexible end. To assess this, we applied our flexible-aware version of the *RBD* score. We observed that *RBD* score improves for all flexible universes (Fig. 7B). The change is less significant for CCF universe; however for LH and HMM universes, the new score is close to one, with the HMM universe performing slightly better.

We computed a version of the likelihood score that considers universe flexibility. This score shows a significant improvement; for all flexible universes, all collection scores change by an order of magnitude (Fig. 7C). Likelihood values that consider flexibility are similar for all universes; however, the HMM universe performs slightly worse for large ChIP-Seq collections (*CTCF ChIP large, TF ChIP*). This reflects that, unlike the CCF and LH universes, the HMM universe does not explicitly optimize likelihood. A more detailed look into likelihood showed that although the HMM universe has the best likelihood of cores of the regions, it performs less well for boundary positions (see Methods; Fig. S6).

### Assessing across metrics

Since each metric assesses different aspects of universe fit, considering them independently limits the scope of assessment. For a holistic view, we summarized the scores of each metric into normalized heatmaps, allowing comparison within and across metrics (Fig. 8). We observed several informative cross-score patterns: First, the *F*_10_-score is the most consistent metric across universes, indicating that all these universes cover the collections to similar extent (Fig. 8A). In contrast, the *RBD* indicates more variation in how well universes represent region boundaries (Fig. 8B). This is consistent with our intuition that matching boundaries is a more difficult task, since it requires ensuring large regions are split well. The disparity between *F*_10_-scores and *RBD* score demonstrates their combined utility. For example, the union universe has perhaps the overall best *F*_10_-scores across universes, but has low *RBD* score. Inversely, the SCREEN universe has the best *RBD* score for the *B-LCL ATAC* collection, but is significantly worse than any data-driven universe for *F*_10_-score. In likelihood scores, the CC, CCF, LH universes outperformed other universes to similar extent, while tiles, RB and union were universally poor fits for all universes (Fig. 8C). The HMM has much better *RBD* score than CC, but a worse likelihood, reflecting that likelihood considers more than boundary positions. Additionally, the likelihood is stricter in boundary assessments: while for *RBD* score we take a median of actual values, for likelihood we use probability of a given position being a boundary. Comparison of flexible and fixed versions of *RBD* score and likelihood highlights the value of flexible regions (Fig. 8D, E); *RBD* score for LH and HMM improved significantly after accounting for flexibility.

**Figure 8:**
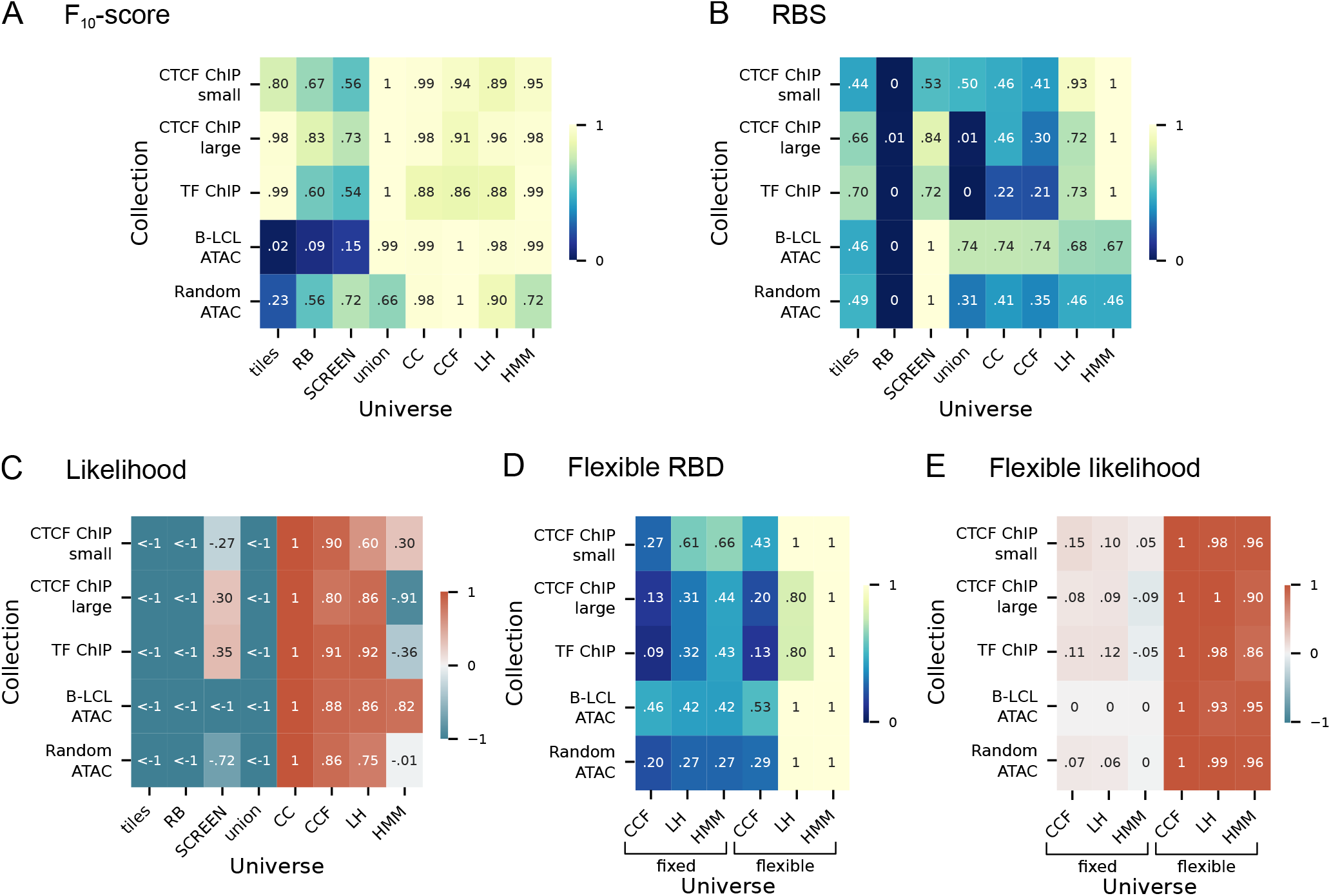
Results of universe comparison using different scores. A) Row normalized F_10_-score of each universe given collection. B) Row normalized RBD score of each universe given collection. C) Row normalized likelihood of each universe given collection. D) Row normalized flexible version of RBD score of each universe given collection. E) Row normalized flexible version of likelihood of each universe given collection.

### Application on downstream analysis

To demonstrate how universe affects downstream analyses and how our universe building and evaluation methods can be applied, we performed a region set enrichment analysis using LOLA, a tool for statistical region enrichment analysis (15). The goal is to take some demo region sets and then use them to search a database of region sets to find similar regions, and explore how the choice of universe affects the results. We constructed two different experiments. For our first experiment, we used the *Random ATAC* collection as a database. We used three predefined universes – tiles, RB and SCREEN – as well as data-driven universes built from the *Random ATAC* and *B-LCL ATAC* collections. To see how the universes based on rare cell types perform, we also added data-driven universes built from fifteen Glia ATAC-Seq files. We queried the database with fifteen files that were not present in the database or were not used for universe construction, representing three different cell types: five files from A549, five from B-LCL, and five from Glia samples. For our second experiment, we used 623 ChIP-Seq files representing different TFs to build a database and data-driven universes. We queried the database with thirty files representing different TFs: ten files from EZH2, ten files from POLR2A, ten files from YY1. For both experiments, we assessed performance of the region enrichment analysis with *R*-precision (rPPV), which measures the precision based on the top *R* results, where *R* is equal to the number of correct results in the whole database.

Our results demonstrate clear impact of the universe on analysis performance (Fig. 9A). In the first experiment, the data-driven universes were the best performers for the specific questions; Glia ATAC queries (gray dots) performed best under the tailored Glia data-driven universes, followed by the *Random ATAC* data-driven universes, and performed poorly with any pre-defined universes or with the B-LCL data-driven universes. The B-LCL queries (yellow dots) also performed best with the tailored B-LCL-driven universes, and performed reasonably well with predifined or *Random ATAC* datadriven universes, but poorly with the Glia data-driven universe. Finally, A549 samples performed equally well with predefined or *Random ATAC* data-driven universes, and poorly with the universes built on the other data types. This shows that using universes based on a specific cell type increase the performance of downstream analysis for this type. Our second experiment, based on TF data, shows a clear difference between the union universe and other more complex data driven universes. We also see an imbalance among the predefined universes, with the SCREEN universe outperforming tiles and RB in general. In conclusion, for ATAC-Seq data, rPPV is higher for data driven universes than predefined ones; for TF data rPPV is higher for more complex data-driven universes (CC, CCF, ML, HMM) than union universes. Overall, these results demonstrate that choosing the right universe is a complex task. In our experiments, the more complex data-driven universes (CC, CCF, LH, and HMM) are always either comparable or superior to simpler data-driven or pre-defined universes, although the exact performance depends on the both the initial data and the downstream task. Most importantly, this analysis suggests that the assessment methods we present can be helpful in choosing the right universe.

**Figure 9:**
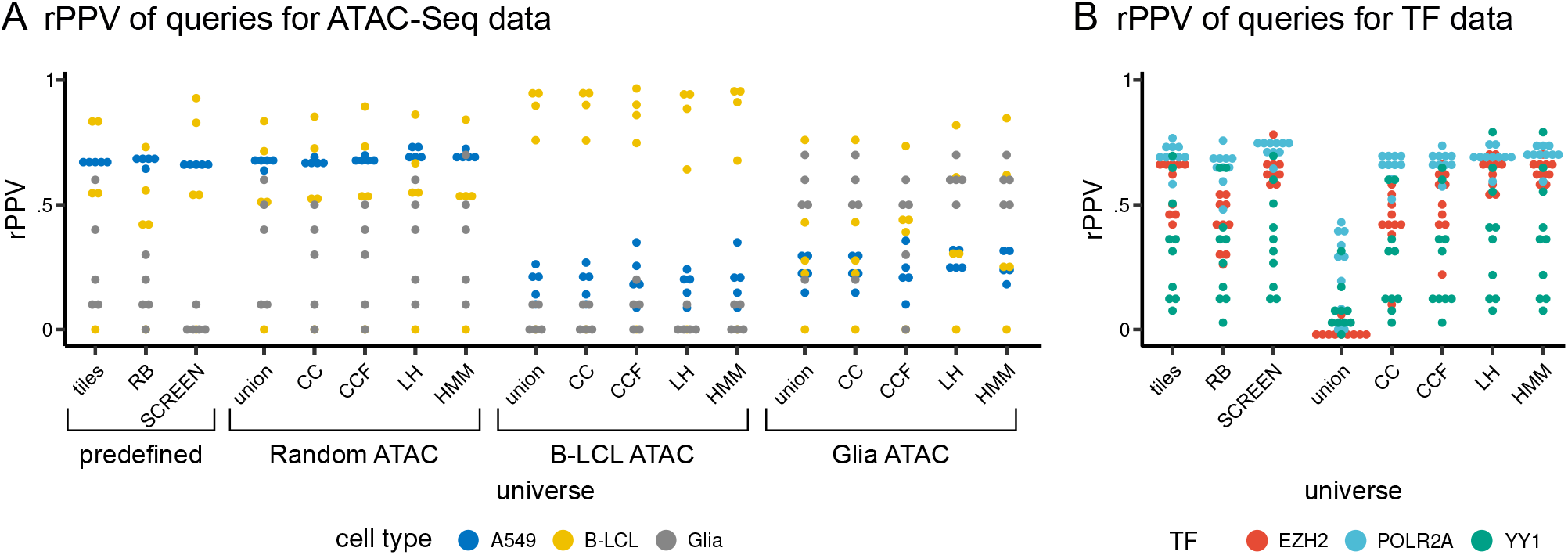
Results of downstream enrichment analysis depends on universe. A) R-precision (rPPV) of query files depending on universe for ATAC-seq experimental set. B) R-precision (rPPV) of query files depending on universe for ChIP-seq experimental set.

## Discussion

Many integrative epigenome analyses require the data to be defined on a set of consensus regions, or universe. This universe is critical for analysis because it determines the precision of regions assessed. Our experiments highlight how different universes can have different levels of fit to collections and may therefore be useful for different tasks. Despite the importance of this choice, few approaches have been developed to aid analysts in building appropriate universes or assessing the fit of an existing universe. In this study, we have addressed these issues by presenting several novel methods to build universes from collections of region sets, as well as new ways to assess the fit of a universe to a collection of region sets. We also introduced the concept of flexible segments, and proposed several methods for constructing universes that can use either traditional fixed boundaries or flexible interval boundaries.

In general, data-driven universes outperformed predefined ones. However, the data-driven universes also have a weakness: by definition, they change with the underlying collection, and therefore cannot support an integrative analysis that spans collections. If results need to be compared across collection, then a shared universe is required. There are a few options for analysis in this case, all of which are facilitated by our work: First, a custom data-driven universe that spans all included collections can be built. Since we have described several ways to create well-performing data-driven universes, it would be easy to just design a bespoke universe for a given comparative analysis. However, this may not always be possible or convenient, and at some point, an external universe may be preferred. In this case, a tradeoff is required: fit of the universe must be sacrificed to increase interoperability with other collections. Our assessment methods now provide a principled way to assess this trade-off and inform research decisions.

Among the non-data-driven universes we tested, SCREEN performed well for the collections we tested. However, there are almost certainly other collections or use cases for which the tiles, RB, or other external universes would be a better choice. Along these lines, we propose that our methods for building universes can be used in the future to create predefined universes from large collections, thereby creating even better global universes that can be re-used for integrative analysis. In the future, a centralized repository of universes, built using different methods and for different target use cases, could be a useful resource; a given collection of region sets could be represented into different universes based on the balance of fit and need for integration.

Based on our results, we propose that the union universe, though widely used, does not represent ChIP-Seq data well, particularly for large collections. Instead, we propose the HMM universe as a good all-around option that solves many of the issues with simpler universes. It has the highest sensitivity to boundary positions and good recall. It also provides adjustable parameters; by setting emission and transition probabilities, users may adjust model sensitivity and keep it consistent across collections. Still, the final choice of universe should consider the needs of downstream analysis. Our results show that assessing universe fit is a complex question, with many features to optimize. Even with our assessment metrics, it is difficult to claim an optimal universe for a given collection; instead, the answer depends on the downstream analysis priorities. For example, in general, the CCF and LH universe represent properties of a whole collection well, but they exclude infrequent regions; thus, they may be useful for NLP analysis of the genome but could lead to losing information about rare cell types in single cell analysis. In contrast, the union universe by definition covers all bases found in the region set collection, and therefore has a great *F*_10_-score; however, it also merges regions, which is reflected by poor *RBD* score and likelihood scores, indicating that it would not be a good fit for an application that requires high region resolution. Therefore, multiple perspectives must be considered for a holistic assessment of universe fit.

One advantage of our new universe construction methods is that they naturally create flexible universes. Flexible universes are a new way of summarizing information from large collections with less loss of information. We showed that proposed approaches of making flexible universes improve results over inflexible universes. We see flexible regions as a powerful new concept that can modify our current way of thinking about universes. Furthermore, we expect that using them for differential peak analysis, statistical region enrichment analysis and NLP approaches has potential to improve results. Flexible regions will become more useful as we and others develop the necessary tooling to work with them; for example, we will require tokenization methods that can project a traditional region set into a flexible universe quickly and accurately, while considering the universe flexibility.

In conclusion, this research provides new concepts, methods, and insight that will help researchers to determine the best analysis path for many types of genomic region analysis.

## Supplementary Methods

### Building input coverage tracks

The input to our universe-building methods is genome signal tracks that count the sum of start positions, end positions, and overlaps at each base. To do this, we developed a fast algorithm that allows us to produce all three of these tracks using a single pass through the reads.

First, each region set in the collection is sorted by sequence name (chromosome) and start position. Then, these region sets are merged into a single, sorted region set that contains all regions in the collection; this combined region set now contains potentially many overlapping regions. We process each chromosome independently. For each chromosome, we first sort the ends separately. This decouples the interval pairs, so they no longer represent regions; for our use cases, we don’t require that pairing, and this step allows us to process the files in a single pass, making the computation very fast.

We initialize a coverage variable to 0, which will indicate the coverage for the current position in the genome. To make signal tracks more stable, for start and end positions, we smooth the tracks by a smoothing window size *w*. To do this, we adjust the boundary position by subtracting *w/*2, and push to a stack of “smoothed window ends” the value of *p* + *w/*2. This effectively turns the start and end position into a region; for starts we have their start and end values, for ends we have their start and end values. They thus can use the same algorithm we use for the coverage tracks, which simply use the original (non-smoothed) start and end positions.

Given a list of sorted starts and ends, we use a triple plane-sweep algorithm to go through the positions in the genome and the sorted starts and ends using nested loops. In the outer loop, we loop through bases in the genome; in the inner loop, through regions. At each position in the genome, we increment *coverage* variable for region starts at this position, and decrement the variable for each interval that ends. We then we emit the current number of overlapping regions. This yields a base-pair-level signal track. The same algorithm can be used for smoothed start, smoothed end, or raw coverage signal tracks.

#### Dealing with noise through smoothing

The point of this smoothing step is to deal with noise in the region boundaries. Experimental sets of regions, like most measurements, contain noise. Addressing this noise is a core part of what the models we are building is to do. An example of a consensus algorithm that does not consider noise is the simple union approach, which will pick up all regions that show up in any input region set. The goal of the more sophisticated models is to reduce noise, without being too aggressive and eliminating true signal. Smoothing deals with noise by accommodating randomness in the exact placement of a peak start and end. Before we feed our observed tracks into the HMM, we smooth the peak boundaries (by a customizable parameter). This has the effect of canceling out variation due to noise in starting location. For practical purposes, the starting position of a peak is often not important down to a specific single nucleotide; hence, smoothing out across a range of 10, 25, or 50 bases allows the model to more accurately hone in on the most likely start and end positions. The level of smoothing can be tuned depending on the level of noise in the input data.

### Coverage cutoff universe

#### Probabilistic models of coverage and background

To improve our description of how a genome is covered by collection of files, we propose a probabilistic approach. Given a collection of *n* region sets ℝ = [ ℛ_1_, ℛ_2_, … ℛ_*n*_] where ℛ_*i*_ denotes a region set ℛ_*i*_ = [*r*_1_, *r*_2_, …*r*_*m*_], and *r*_*i*_ denotes a single region in the set, we make a model describing the probability of each position in the genome being covered by a region in a randomly selected region set from the collection. To compute this probability, we first compute the coverage frequency for each genomic position independently, *freq*_*c*_, a vector of length *g* (the length of the genome). We do this by simply counting, for each position, in how many files it is covered, yielding frequencies of coverage for each genomic position. We then convert *freq*_*c*_ into a probability distribution over the genome by dividing each element by the sum. We define the probability of a position being covered as:

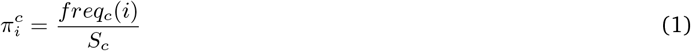

where *i* indicates the position (base-pair) and *S*_*c*_ denotes sum of coverage frequency counts across the whole genome:

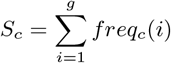

This variable, 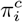, can be thought of as an extension of a multinomial random variable. Consider a multinomial random variable, where the categories are positions (individual base pairs) in the genome. We seek to define a distribution across these positions. However, our model does not follow a simple sampling model either with replacement or without replacement; instead, we employ a hierarchical sampling model, where samples (base pairs) are first grouped (representing files), where groups may not duplicate items, indicating a without replacement sampling strategy. But across groups, duplicates are allowed, indicating a with replacement sampling strategy. Thus, 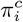 represents a probability distribution across the genome of whether a base is covered in the region set collection ℝ. In other words, if we think of drawing random bases from a set of files, this distribution describes the probability of selecting each base, using our hierarchical sampling strategy.

Next, we calculate an independent probability distribution for the inverse model, the probability of each position being *not covered* in our sample data, which we call background:

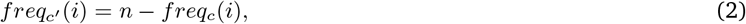

where *i* again represents genome position and *n* is the number of files. As before, we divide these frequencies by their sum to get a probability distribution across the genome, this time of background probabilities:

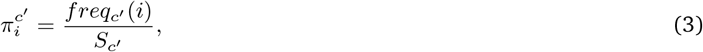

where *S*_*c*_ ′ denotes sum of genome background (the total number of bases that were not covered in a set of files):

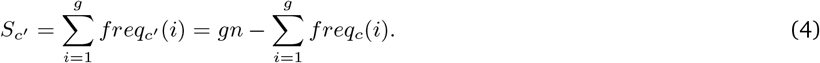

Where *g* is the length of the genome, Thus, *gn* represents the maximum possible elements that could be covered in a given collection of *n* files; the unit could be “file-bases”.

Combing information from those two probabilities, *π*_*c*_′ and *π*_*c*_, we get a model *M* describing for each position how probable it is for it to be covered (core) or not (background).

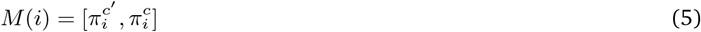

These two probability distributions are not entirely independent, but they are also not exact inverses because of the complicated file structure, which affects the sampling strategy.

#### Finding likelihood cut-off value

We can use presented model to find the most probable universe (under the model). To do it, we have identify all the bases for which the probability of being covered is higher than the probability of being not covered; that is, those that satisfy the following inequality:

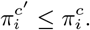

Substituting the definitions from (1) and (3), this is equivalent to:

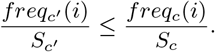

Then, usingequations (2) and (4), we get:

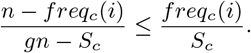

Multiply both sides by *S*_*c*_(*gn* − *S*_*c*_):

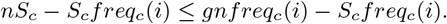

Simplifying:

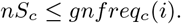

Solving this equation for *freq*_*c*_(*i*), results in:

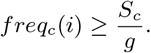

Thus, if we have identified a cut-off value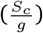; we take the maximum likelihood universe under this model for a given collection as the one for which any locations *freq*_*c*_(*i*) above the cutoff are included, and others are excluded.

### Maximum likelihood universe

#### Probabilistic model extended to region boundaries

The previous probabilistic description a region set collection across genome considers only region coverage; it does not include information about the position of regions’ boundaries. To extend the model, we make similar models for start and end boundary probabilities. We calculate the frequencies of each position containing either start *freq*_*s*_ or end *freq*_*e*_. Moreover, as before, for each base pair, we count the number of region sets in which it is background (in other words, *not* a start), for start as *freq*_*s*_′ = *n* − *freq*_*s*_, and for end (*not* an end) as *freq*_*e*_′ = *n* − *freq*_*e*_. Using that, we derive for each position the probability of it being a boundary or boundary’s background:

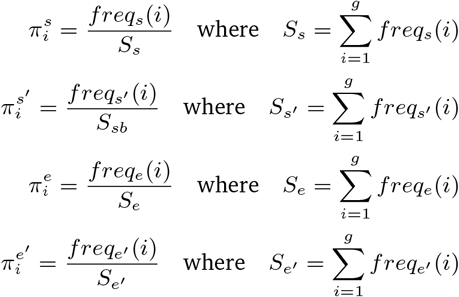

Thus, 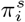 is the probability of being a start, 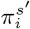 is the probabilty of *not* being a start, 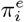 is the probability of being an end, and 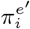 is the probabilty of *not* being an end.

Combining these probabilities with the model described in (5) we get a complex model describing the probability distribution of a collection of files across the genome:

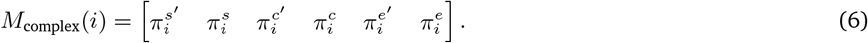

#### Building maximum likelihood universe

We can use this extended likelihood model (6) to derive a maximum likelihood flexible universe. We do it by finding an optimal genome segmentation with four states: start *s*, core *c*, end *e*, and background *b*. These states reflect the parts of a flexible region. We assume that this can be found by dividing the problem into overlapping sub-problems. That means that the best segmentation of first *i* positions is a result of the most likely segmentation of first *i* − 1 positions and the most likely state at position *i*. This assumption makes the problem amenable to dynamic programming, which we employ to find the maximum likelihood universe.

Next, we need to create a scoring function that considers all the probabilities described earlier. We reason that a score for assigning a particular base to a particular state could be computed by multiplying the probability of the state with the probabilities of *not* being in the other states. For example, we compute the probability of being in state *s* as 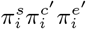, and equivalently for state *c* and state *e*. To compute the probability of being in none of these states, or, equivalently, in the overall background state *b*, we can compute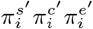 (multiplying all three background probabilities). Using this approach, we create a scoring function *P* that for each position in the genome *i* denotes its probability of being in a given state:

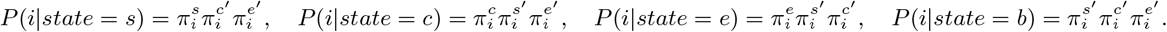

Using *P*, we can represent our problem as a recursive equation:

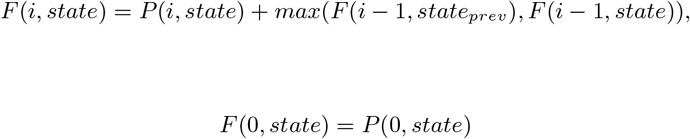

where *i* denotes position in the genome, *state* ∈ {*s, c, e, b*} is the current state, and *state*_prev_ a state from which *state* can be accessed, for example if *state* = *s* than *state*_prev_ ∈ {*s, b*}. Using this equation, we use dynamic programming to find a most likely path.

#### LH universe filtering

Although the LH universe is designed to find an optimal universe, it can result in very small universe regions. We remedy that by removing small regions (below 100 bp). Furthermore, for flexible likelihood universes, we also add a filtering step; for each region we calculate its contribution to the flexible universe likelihood by calculating the difference between the total universe likelihood and the likelihood with the current region removed (Fig. S2). This *flexible universe likelihood* score is based on a different conceptualization of the region likelihoods (see section on universe likelihood in Methods), and thus, individual regions may contribute negatively to that score. Based on that, we filter out regions with negative contribution to the universe likelihood.

### Hidden Markov Model universe

#### Process of determining HMM parameters

One advantage of the HMM is that we have control over parameters for the transition and emission probabilities, allowing us to tweak the model. In many applications of HMMs, these parameters are trained using gold-standard reference input data and the forward-backward algorithm. However, in our case, we lack suitable training data. We attempted to construct reference datasets that could be used to train parameters, but were unsatisfied with the final trained models, which would often tend to converge to parameter sets that yielded no universe regions. Instead, we decided to parameterize the HMM manually by tuning the system to yield the type of results we sought. We found that, through trial and error, we were able to achieve better results.

In the process of tuning the HMM and applying it to different sizes of input region set collections, we realized that the parameters would need to vary depending on the input dataset. Since we desired a universal model that could be applied in many different scenarios, we reasoned that a model based on a pre-processed normalized signal could be made to work across varying input scenarios. Therefore, we introduced a pre-processing step of quantile normalization of the input signal tracks. For each track, we map its values to a reference distribution, in a way that normalizes the distribution to have the same quantile values as the reference distribution. As a reference, we use Negative Binomial distribution, since in most cases it represents genome coverage by the collection, with different success probabilities for cores and boundaries, respectively 0.2 and 0.1, and same number of successes, which is equal to 1. Then, we feed the normalized signals into the HMM. This approach allows us to use a single set of emission probabilities for diverse collection sizes, thereby enabling the HMM to process diverse data collections without requiring collection-specific tuning.

This quantile normalization step allows us to handle different sizes of input collection, making it more universally useful; however, it may still be desirable to tune the model differently depending on the noise of the input collection. Tuning the emission and transition probabilities also allows us to dial the sensitivity and specificity of the model, which provides the ability to handle data with different single-to-noise ratios. For example, by allowing the background state to emit higher start, coverage, and end signal, the model will be less likely to create universe regions where a small minority of input sets are covered, and vice versa. The final model we used is based on one empirical evaluation of universes and attempt to strike a practical balance, but interested users could tweak the parameters to dial up or down the noise tolerance.

#### Parameters

Since it is very unlikely for the process to start in something different from background, we assign to it the highest probability *p*_start_. In transition matrix *T*, we define the peak structure by setting to zero impossible transitions. Moreover, we set a probability of staying in a given state much higher than probability of transiting to the next. This way, we can make sure that segments are not too short. The third matrix is an emission matrix *E*, which describes probability of given values being emitted from a given state. We used a Poisson distribution with manually tuned parameter *λ* for each emitted variable (starts, coverage, and ends).

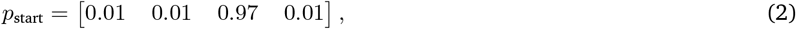

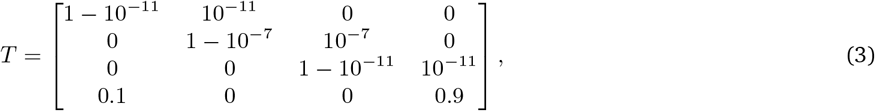

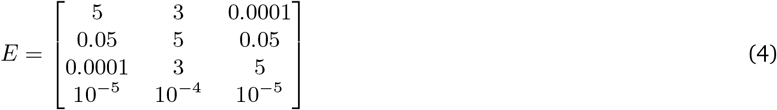

In matrix *T*, the rows and columns correspond to the four states in this order: start, coverage, end, background.

In matrix *E*, the rows are the same: start, coverage, end, and background, and the columns correspond to the 3 observed emission variables: start, coverage, and end.

#### HMM universe filtering

We remove from the raw HMM universe small regions below 100 bp. Additionally, as for the LH universe, we filter regions, which have strong negative contribution to the universe likelihood score (*<*-100) (Fig. S3).

### Calculating flexible universe likelihood given model

#### Overview

We seek to compute a likelihood score for a proposed universe given a collection of region sets. We previously described a likelihood model used to *create* a universe given a collection (which we called the likelihood universe). At first, we tried to apply the original model as a universe assessment method; however, this likelihood model could not be used on flexible universes because boundaries, which are now regions instead of points, contribute very negatively to the score, since the probabilities of emitting a boundary are lower than coverage due to their sparsity. Therefore, we also derived a separate likelihood approach to be used for evaluating flexible universes.

#### Hard universe

First, we describe the simpler case of a hard (not flexible) universe, and we will then extend this to the case of a flexible universe. For hard universes, we *can* use the original likelihood to evaluate, because they don’t have the problem introduced by flexible starts and ends. To calculate the likelihood of any universe given region set collection ℝ, we first create a binary matrix representation of the universe, *U*_*binary*_. We build *U*_*binary*_ with 6 rows, corresponding to 1) start-background, 2) start, 3) core-background, 4) core, 5) end-background, and 6) end. The columns correspond to genome positions. We set the value to one if, at the given position, the universe contains the corresponding part of the peak.

For example, for a region *r* with start *s* = 3 and, end *e* = 7 we represent part of the genome as:

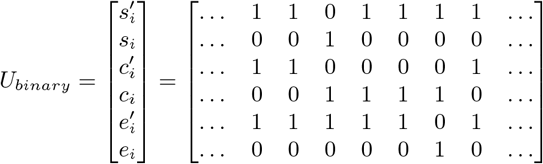

Where *i* corresponds to genome position, yielding a long 6 × *n* matrix, *n* is the length of the sequence.

Then, we can calculate the log-likelihood of the universe using a matrix multiplication:

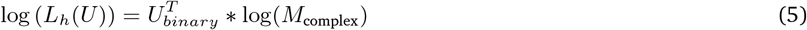

Where *U*^*T*^ indicates the transpose operation.

#### Flexible universe

The flexible universe is trickier because the original data have point boundaries, but the universe has flexible boundaries. Therefore, we cannot compute probabilities in the same way, using the frequency of boundaries in the region set collection.

To address this, we start with a similar idea to calculate likelihood of a flexible universe, but we model start and end not as points, but subregions with uniform probability of containing a given boundary. To accommodate that, we introduce weights in *U*_*binary*_ matrix representation. For example, for region *r* with *s*_start_ = 5, *s*_end_ = 8, *e*_start_ = 15, *e*_end_ = 20 we represent it as:

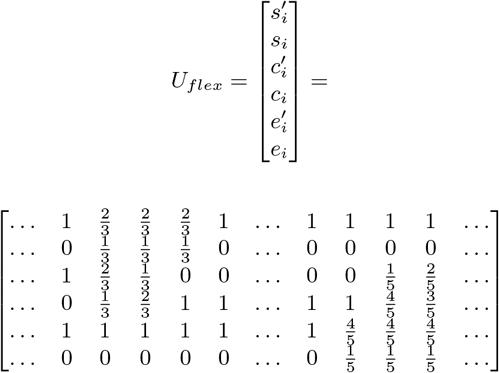

Then, we simply use the same matrix multiplication to calculate the likelihood with this weighted matrix:

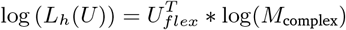

The intuition behind this approach is that the flexible region is not defining a specific point for the region boundary; therefore, the contribution of the likelihood from the boundary distribution should be distributed across the flexible component. For the start and end probabilities, we thus divide the weight evenly across the segment; the denominator thus equals the length of the segment. The corresponding background is set to 1 - the state weight. To accommodate this dispersion of probability score, we also adjust the weights of the core state, which increases as the genome progresses because if a start had happened in the preceding base, then the next base would increase in probability of being in a core region. In other words, within a start state, as you progress along the genome, the probability of being in a core should increase. To model this, we simply have the probabilities of being in a core start at zero at the leftmost boundary of the start state, and increase to 1 at the rightmost boundary of the start state. Then, we do the inverse at the end states, starting at 1 and ending at 0.

### General comparison of universe statistics

Data-driven universes tend to cover a similar amount of the genome as the data used to build it, except for universes derived from likelihood models (CC, CCF, LH universes) for large collections (CTCF ChIP large and TF ChIP), where the coverage drops from around 90% for original data to below 50% for universe (Fig. S4). This observation is the consequence of Eq. 1, which states that as the sum of coverage of the genome by collection increases, so does the maximum likelihood cutoff value. That means that for collection with high sum of genome coverage by collections, many positions with low coverage will be excluded. Moreover, union universe produces large peaks, especially for ChIP-Seq collections (CTCF ChIP small, CTCF ChIP large, TF ChIP); for TF collection average peak size is over 40 500, where peak size in the data is around 400. Interestingly, for CTCF ChIP large both HMM and LH universe have many small regions, which reflects the complicated nature of the data.

### Accounting for noise in universe evaluation

The evaluation metrics account for noise through their asymmetry. Many of the evaluation metrics we proposed are asymmetric versions of evaluation parameters; for example, the *F*_10_ score penalizes one direction 10 times more than the other. The point of the asymmetry is that for these models, it is more problematic to miss real regions than to have a few extra regions. This framework allows the level asymmetry to be tuned; so a user who needed a universe that is highly sensitive could dial the metrics in a way to penalize losing information more. This would create universes that have more peaks, but may have more noise. On the other hand, for an application in which it’s important to filter out more noise, the evaluations could be tuned in the other direction.

## Supplementary Tables

**Table S1:** Statistics of demo region set collections. Five collections of regions ranging from 40 to >8,000 files, giving a diverse set of possible use cases where generating a set of consensus regions could help with data integration.

| Collection | N files | Region number | Genome covered (%) | Average region size |
| --- | --- | --- | --- | --- |
| CTCF ChIP small | 40 | 7,107,842 | 42.5% | 367 |
| CTCF ChIP large | 877 | 152,694,264 | 89% | 358 |
| TF ChIP | 8,503 | 1,463,013,952 | 90.7% | 394 |
| B-LCL ATAC | 400 | 776,673 | 0.2 % | 268 |
| Random ATAC | 5,000 | 76,332,694 | 17.8% | 290 |

**Table S2:** Statistics of universes. Basic statistics of all presented universes, including the number of regions in the universe, their average size, and the percentage of the genome they cover.

| Collection | Universe | Region number | Average region size | Genome covered (%) |
| --- | --- | --- | --- | --- |
| CTCF ChIP small | union | 2,320,577 | 566 | 40.9 % |
| CTCF ChIP large | union | 151,846 | 18,116 | 85.7 % |
| TF ChIP | union | 69,209 | 40569 | 87.5 % |
| B-LCL ATAC | union | 14,101 | 357 | 0.2 % |
| Random ATAC | union | 1,306,508 | 421 | 17.1 % |
| CTCF ChIP small | coverage | 2,120,192 | 572 | 37.8 % |
| CTCF ChIP large | coverage | 1,368,624 | 504 | 21.5 % |
| TF ChIP | coverage | 957,516 | 1030 | 30.7 % |
| B-LCL ATAC | coverage | 14,027 | 357 | 0.2 % |
| Random ATAC | coverage | 547,740 | 387 | 6.6 % |
| CTCF ChIP small | CCF | 1,509,661 | 605 | 28.4 % |
| CTCF ChIP large | CCF | 492,846 | 961 | 14.8 % |
| TF ChIP | CCF | 612,181 | 1639 | 31.3 % |
| B-LCL ATAC | CCF | 9,835 | 413 | 0.1 % |
| Random ATAC | CCF | 316,089 | 618 | 6.1 % |
| CTCF ChIP small | ML | 3,381,856 | 264 | 27.8 % |
| CTCF ChIP large | ML | 3,042,529 | 218 | 20.7 % |
| TF ChIP | ML | 2,887,104 | 364 | 32.7 % |
| B-LCL ATAC | ML | 15,538 | 353 | 0.2 % |
| Random ATAC | ML | 694,194 | 382 | 8.3 % |
| CTCF ChIP small | HMM | 4,950,358 | 271 | 41.8 % |
| CTCF ChIP large | HMM | 10,618,927 | 194 | 64.4 % |
| TF ChIP | HMM | 7,896,890 | 308 | 75.7 % |
| B-LCL ATAC | HMM | 11,818 | 406 | 0.1 % |
| Random ATAC | HMM | 1,292,045 | 338 | 13.6 % |
| predefined | tiles1000 | 2,938,211 | 1000 | 91.5 % |
| predefined | Regulatory Build | 548,310 | 950 | 16.2 % |
| predefined | SCREEN | 1,063,878 | 273 | 9.0 % |

## Supplementary Figures

**Figure S1:**
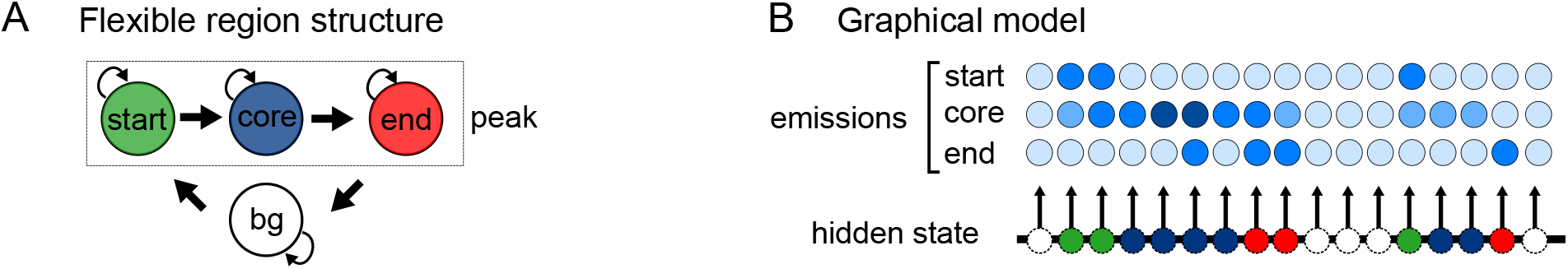
Details of Hidden Markov Model. A) State space diagram showing state transitions for the Hidden Markov Model. This is a graph representation of a flexible region. B) Graphical model describing the sequential states of the HMM.

**Figure S2:**
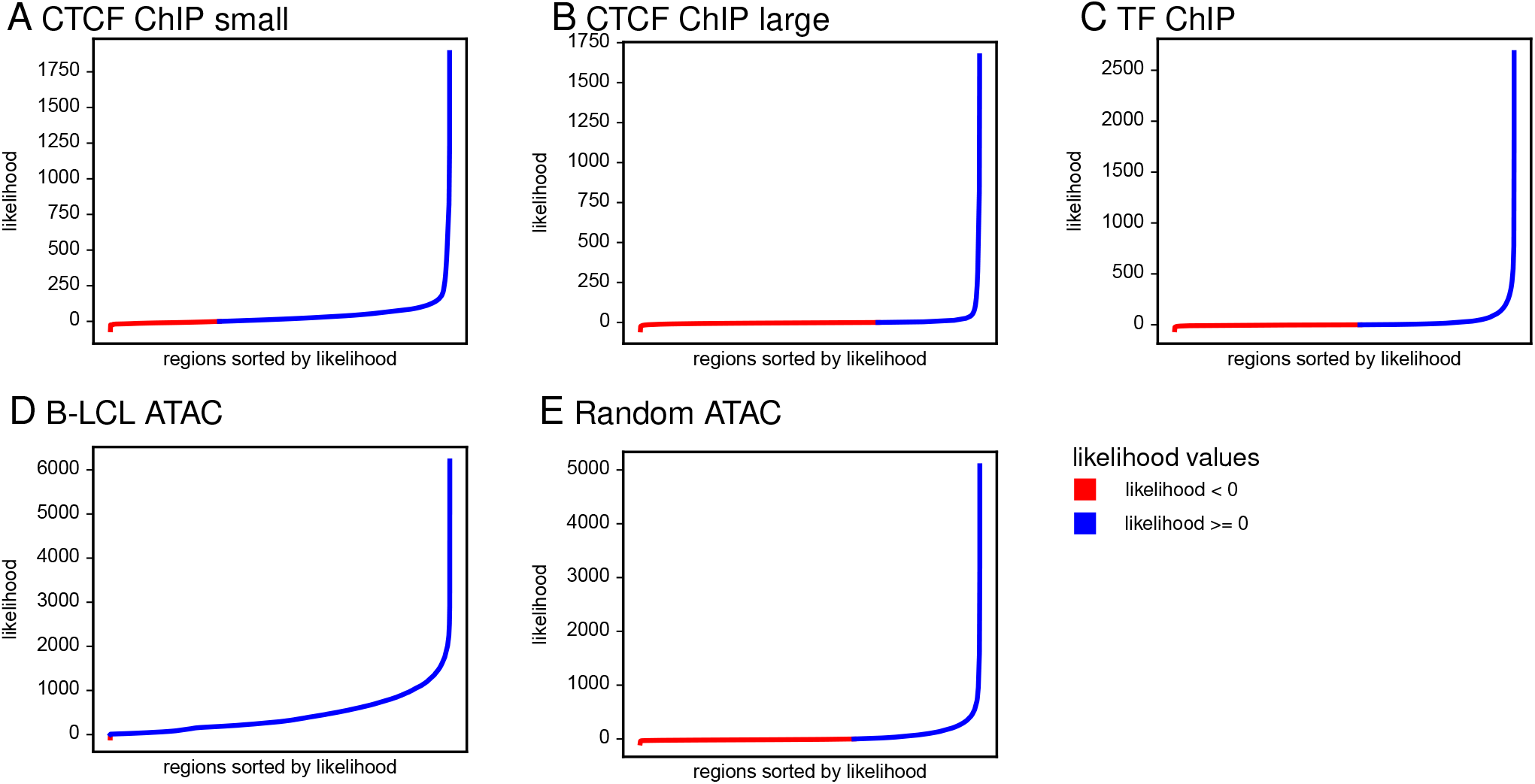
Sorted likelihood inputs of all regions present in the ML universe. A) Likelihoods of regions in ML universe for CTCF ChIP small collection. B) Likelihoods of regions in ML universe for CTCF ChIP large collection. C) Likelihoods of regions in ML universe for TF ChIP collection. D) Likelihoods of regions in ML universe for B-LCL ATAC collection. E) Likelihoods of regions in ML universe for Random ATAC collection.

**Figure S3:**
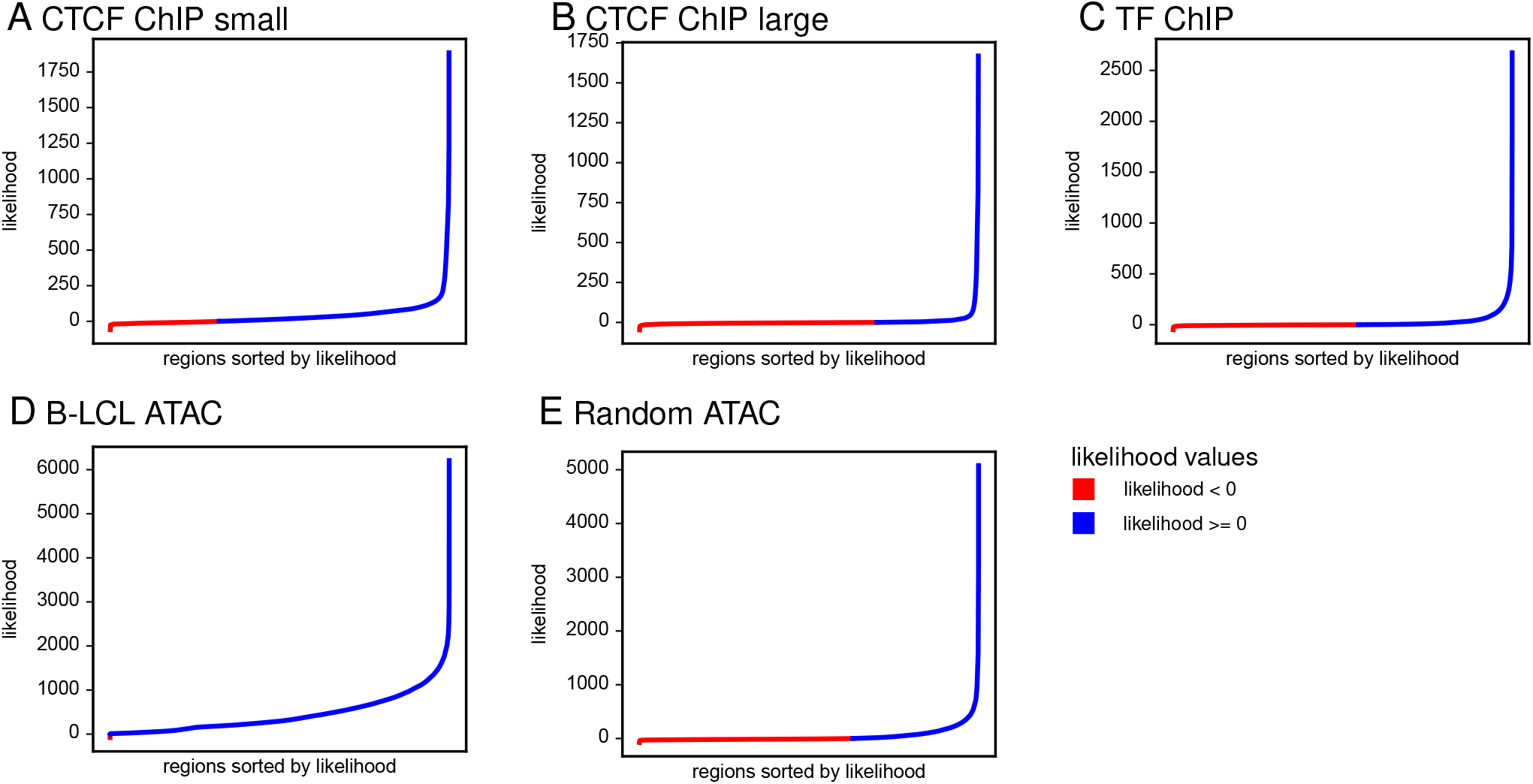
Sorted likelihood inputs of all regions present in the HMM universe. A) Likelihoods of regions in HMM universe for CTCF ChIP small collection. B) Likelihoods of regions in HMM universe for CTCF ChIP large collection. C) Likelihoods of regions in HMM universe for TF ChIP collection. D) Likelihoods of regions in HMM universe for B-LCL ATAC collection. E) Likelihoods of regions in HMM universe for Random ATAC collection.

**Figure S4:**
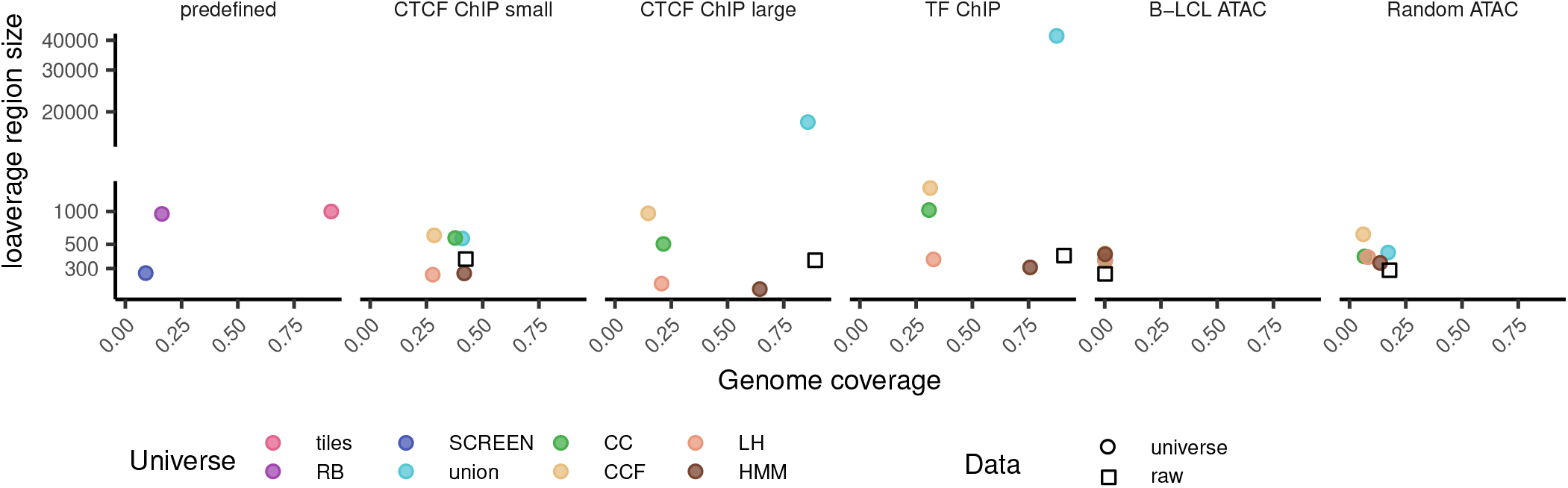
Comparison of universe general statistics. Squares denote properties of raw underlying collection.

**Figure S5:**
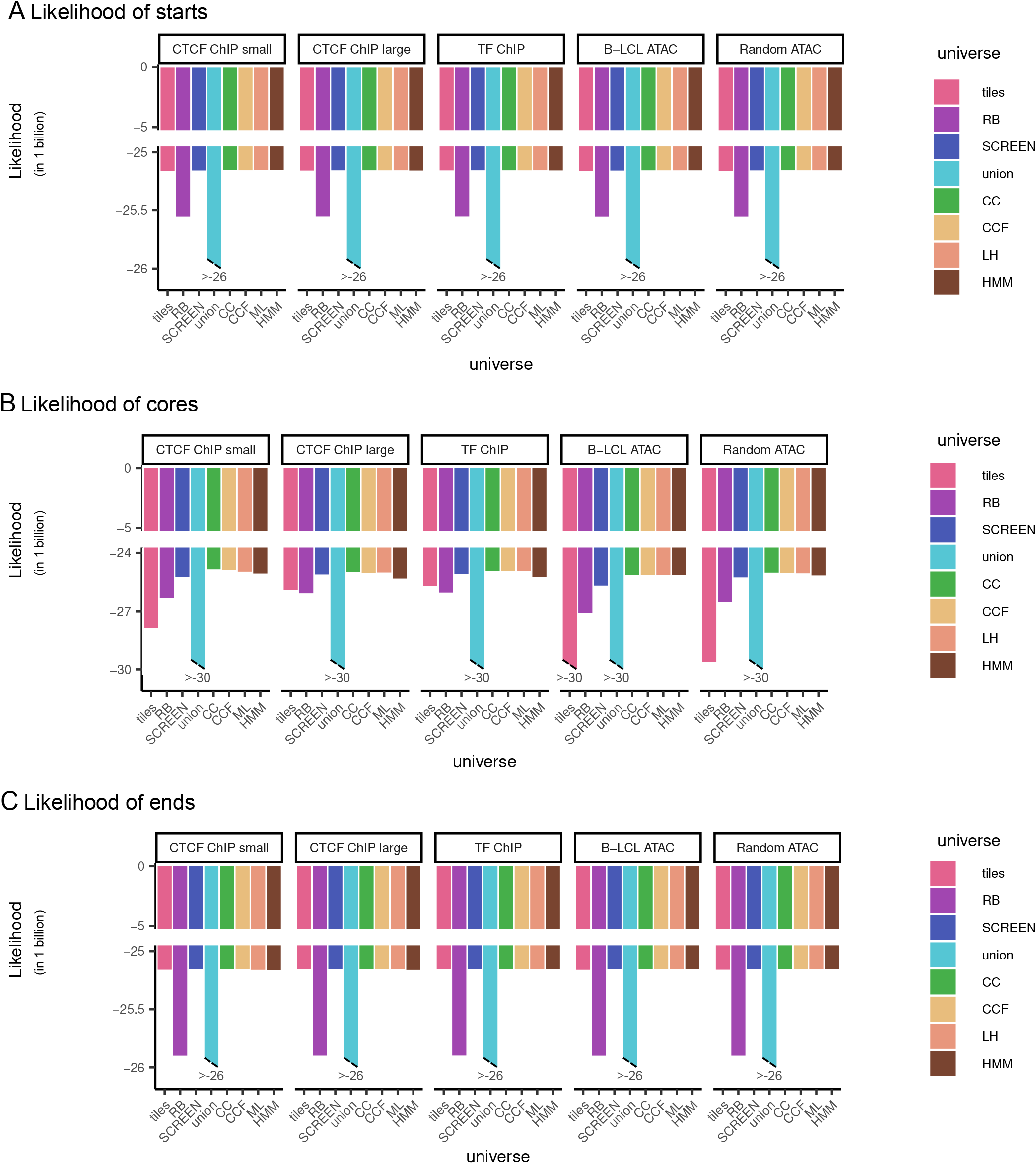
Likelihood of given parts of region. A) Likelihood of region starts by universe and collection. B) Likelihood of region cores by universe and collection. C) Likelihood of region ends by universe and collection.

**Figure S6:**
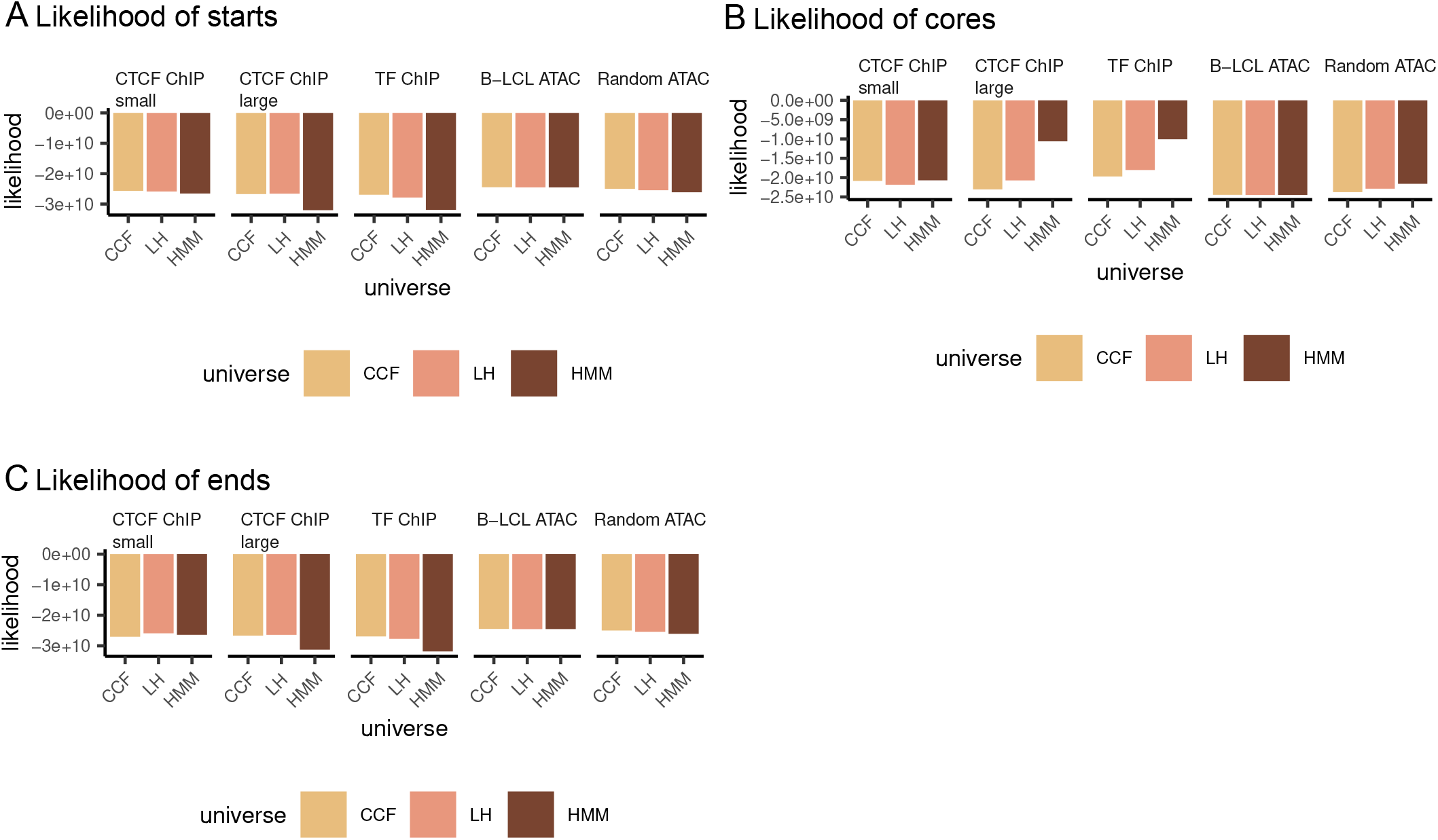
Flexible version of likelihood of given parts of region. A) Flexible likelihood of region starts by universe and collection. B) Flexible likelihood of region cores by universe and collection. C) Flexible likelihood of region ends by universe and collection.

